# Network-based Near-Scalp Personalized Brain Stimulation Targets

**DOI:** 10.1101/2025.05.15.654391

**Authors:** Ru Kong, Aihuiping Xue, Jingwen Cheng, Leon Qi Rong Ooi, Christopher L. Asplund, Xiao Wei Tan, Shih Ee Goh, Jonathan Jie Lee, Jovi Zheng Jie Koh, Rachel Si Yun Tan, Hasvinjit Kaur Gulwant Singh, Trevor Wei Kiat Tan, Alvin P.H. Wong, Ryan D Webler, Michael D Fox, Shan Siddiqi, Phern-Chern Tor, B.T. Thomas Yeo

**Affiliations:** Centre for Sleep and Cognition & Centre for Translational MR Research, Yong Loo Lin School of Medicine, National University of Singapore, Singapore; Department of Electrical and Computer Engineering, National University of Singapore, Singapore; Department of Medicine, Healthy Longevity Translational Research Programme, Human Potential Translational Research Programme & Institute for Digital Medicine (WisDM), Yong Loo Lin School of Medicine, National University of Singapore, Singapore; N.1 Institute for Health, National University of Singapore, Singapore; Integrative Sciences and Engineering Programme (ISEP), National University of Singapore, Singapore; Department of Biomedical Engineering, College of Design and Engineering, National University of Singapore, Singapore, Singapore; Division of Social Sciences, Yale-NUS College, National University of Singapore, Singapore; Department of Psychology, College of Humanities and Sciences, National University of Singapore, Singapore; Institute of Mental Health, Singapore; School of Psychology, Faculty of Health, Medicine and Behavioural Sciences, The University of Queensland, St Lucia, QLD, Australia; Department of Psychiatry, Mass General Brigham, Harvard Medical School, Boston, USA; Center for Brain Circuit Therapeutics, Brigham & Women’s Hospital, Harvard Medical School, Boston, USA; National University Hospital, Singapore; Martinos Center for Biomedical Imaging, Massachusetts General Hospital, Charlestown, MA, USA

## Abstract

Functional connectivity (FC) is often used to identify personalized targets for transcranial magnetic stimulation (TMS). However, existing methods often overlook individual differences in whole-cortex network organization. Furthermore, in some personalized TMS protocols, lower stimulation intensity is used for targets closer to the scalp, which may improve patient tolerance. Here, we develop an algorithm to simultaneously optimize FC and scalp proximity for target localization. We first use the multi-session hierarchical Bayesian model (MS-HBM) to estimate high-quality individual-specific cortical networks. A tree-based algorithm is then used to select the optimal target. With essentially no parameter to tune, our framework may potentially improve generalizability across populations. We compare our approach to existing “cluster” and “cone” algorithms. In two test-retest datasets of healthy individuals from the United States and Singapore, tree-based MS-HBM reliably identifies personalized TMS targets for depression near the scalp. Tree-based MS-HBM targets compare favorably with cluster and cone targets in terms of reliability, scalp proximity, and FC to the subgenual anterior cingulate cortex (sACC) in new out-of-sample MRI sessions. To demonstrate versatility, the same algorithm identifies personalized anxiety targets without tuning any parameter. In patients with treatment-resistant depression, tree-based MS-HBM targets compare favorably with cluster and cone targets in terms of reliability, scalp proximity, and sACC FC, hypothetically reducing stimulation intensity by 15% and 5% respectively. MS-HBM also exhibits the best (most negative) electric-field hotspot sACC FC and highest reliability in induced electric fields. Overall, tree-based MS-HBM provides a robust, generalizable framework to estimate near-scalp personalized targets across populations.

## Introduction

Brain stimulation has emerged as a powerful alternative for patients with psychiatric conditions resistant to conventional treatments, including major depressive disorder and obsessive-compulsive disorder (George et al., 1995; Berlim et al., 2013). In depression, 30% of patients fail to respond to any medication or psychotherapy (Sinyor et al., 2010). Consequently, transcranial magnetic stimulation (TMS) targeting the dorsolateral prefrontal cortex (DLPFC) has been approved by the U.S. Food and Drug Administration (FDA) for treatment-resistant depression (Padberg and George, 2009; George et al., 2010). However, TMS response rates are modest (Liu et al., 2014; Sehatzadeh et al., 2019; Sackeim et al., 2020; Ye et al., 2024; Goodman et al., 2025).

One possible reason for the modest TMS response is the reliance on anatomically derived targets, either through scalp-based measurements (Beam et al., 2009) or anatomical registration to a group-level MRI template (Blumberger et al., 2018). These approaches do not account for inter-individual differences in functional organization beyond anatomical variability (Mueller et al., 2013; Laumann et al., 2015; Lynch et al., 2024). However, such differences in functional organization may affect outcomes. For example, patients whose stimulation sites were closer to DLPFC regions, which exhibited more negative resting-state functional connectivity (FC) with the subgenual anterior cingulate cortex (sACC), experienced better treatment outcomes (Cash et al., 2021a; Siddiqi et al., 2021; Kong et al., 2022; Stöhrmann et al., 2023; Cash and Zalesky, 2024; Chen et al., 2025). Several prospective trials have reported impressive outcomes with the sACC-FC target (Siddiqi et al., 2019; Cole et al., 2020; Li et al., 2024; Hearne et al., 2025; but see DeSouza et al., 2024), fueling enthusiasm for personalized connectome-guided targeting, potentially beyond depression.

Furthermore, psychiatric disorders are increasingly recognized as network-level disruptions, rather than dysfunctions of isolated brain regions (Fox and Greicius, 2010; Menon, 2011; Sha et al., 2019). While current brain stimulation practices often target single regions, their therapeutic effects likely arise from modulating distributed brain networks (Eldaief et al., 2011; Liston et al., 2014; Cocchi et al., 2016; He et al., 2024). However, the potential of network-based approaches to personalize stimulation targets remains under-explored (Siddiqi et al., 2023). Recent advances in MRI acquisition (Setsompop et al., 2012; Lynch et al., 2020) and algorithms (Kong et al., 2019; Hermosillo et al., 2024) now enable high-quality, individual-specific network estimation using resting-state fMRI (rs-fMRI). For example, with just 10-min fMRI scans, the multi-session hierarchical Bayesian model (MS-HBM) generates cortical networks of equivalent quality to other approaches using 50 mins of fMRI (Kong et al., 2019). MS-HBM network topography also aligns with task-evoked fMRI activity (Du et al., 2024), predicts cognition and mental health (Kong et al., 2019), and is heritable (Anderson et al., 2021). Selecting personalized targets based on MS-HBM networks might therefore improve target quality.

Another consideration for TMS target selection is the distance between the target and the scalp. In the case of the seminal SNT (Stanford accelerated intelligent neuromodulation therapy) protocol, TMS stimulation intensity is titrated based on the target’s distance from the scalp, i.e., every millimeter reduction in distance-to-scalp translates to 2.7% reduction in TMS intensity (Stokes et al., 2005; Cole et al., 2020, 2022). Stimulation intensity decays with distance from the TMS device, so the assumption here is that deeper targets require stronger stimulation intensity to maintain the same effects (Cole et al., 2020, 2022). Furthermore, common side effects of TMS include headaches. Lower stimulation intensity might potentially enhance patient tolerability, so targets closer to the scalp might be desirable. Some algorithms for TMS target selection select for scalp proximity, e.g., “cone” algorithm (Fox et al., 2013). However, other approaches do not account for scalp proximity, e.g., “cluster” algorithm (Cash et al., 2021b).

Here, we present a network-based algorithm to select personalized TMS targets that explicitly optimize both FC and scalp proximity. We first use MS-HBM to derive individualized cortical networks. A novel tree-based algorithm then aggregates information across varying FC and scalp proximity thresholds to pinpoint an optimal stimulation site. By adapting to individual variability rather than relying on fixed thresholds, our approach has essentially no “free” parameter, thus potentially enhancing generalizability across datasets and psychiatric conditions. In two independent healthy datasets, tree-based MS-HBM identifies personalized depression targets significantly closer to the scalp than the “cluster” algorithm (Cash et al., 2021b), which seeks reliable targets with strong negative FC to sACC without considering scalp proximity. Compared with cluster (Cash et al., 2021b) and cone (Fox et al., 2013) algorithms, tree-based MS-HBM targets exhibit more negative sACC FC. To illustrate the flexibility of our approach, we apply tree-based MS-HBM – without tuning any parameter – to identify anxiety targets based on the anxiosomatic circuit map (Siddiqi et al., 2020), which again compare favorably with cluster and cone targets. Finally, in individuals with treatment-resistant depression, tree-based MS-HBM targets show the best scalp proximity, most negative sACC FC, most negative electric-field hotspot sACC FC, and highest electric-field reliability. Collectively, these results highlight the adaptability of tree-based MS-HBM for deriving personalized stimulation targets.

## Methods

### Datasets

We first considered two test-retest datasets with healthy participants: the Human Connectome Project (HCP) test-retest dataset (Van Essen et al., 2012a; Smith et al., 2013) and a test-retest dataset from Singapore (SING). Both datasets contained structural MRI and resting-state fMRI (rs-fMRI). The HCP test-retest dataset comprised 46 participants with 4 sessions of rs-fMRI. Two sessions were collected on two consecutive days followed by two more sessions (also on two consecutive days) on average 11.6 months after the first two sessions. Each session contained two rs-fMRI runs, each of length 14 min 33 sec. HCP is a publicly available dataset. Use of the HCP dataset is approved by the National University of Singapore (NUS) institutional review board (IRB).

The SING test-retest dataset comprised 18 participants with 4 sessions of rs-fMRI. The first two sessions were collected in two consecutive days, while the last two sessions were collected in another two consecutive days 2 weeks after the first two sessions. Each session contained a single 10-min rs-fMRI run. Participants provided written consent, and data collection was approved by the NUS IRB.

In addition, we included a dataset (SING-D), comprising 43 patients with treatment-resistant depression. Each participant underwent two 10-min rs-fMRI runs acquired in a single MRI session. All data were collected with appropriate ethical approval (DSRB 2023/000397 and 2023/00680).

After pre-processing and quality control of the HCP, SING and SING-D datasets using a pipeline established in our previous studies (Li et al., 2019; Kong et al., 2021), we considered a subset of participants with all rs-fMRI runs. Therefore, our main analyses comprised 32 HCP participants (23 females, 9 males), 18 SING participants (12 females and 6 males), and 43 SING-D participants (20 females and 23 males).

### Acquisition parameters

The HCP acquisition parameters can be found elsewhere (Van Essen et al., 2012a; Smith et al., 2013). Briefly, all imaging data were collected on a customized 3T Siemens Skyra 3T scanner at Washington University in St. Louis. The structural images were obtained with T1w MPRAGE sequence using the following acquision parameters: TR = 2400 ms; TE = 2.14 ms; FOV = 224 × 224 mm; voxel resolution = 0.7 × 0.7 × 0.7-mm voxels. The rs-fMRI data were obatained with the following parameters: TR = 720 ms; TE = 33.1 ms; multiband factor 8; flip angle = 52°; FOV = 208 × 180 mm; voxel resolution = 2.0 × 2.0 × 2.0-mm voxels.

In the case of the SING and SING-D datasets, the acquisition parameters are as follows. All imaging data were collected on a Siemens Prisma 3T scanner at National University of Singapore. The structural data were obtained with T1w MPRAGE sequence using the following acquisition parameters: TR = 2200 ms; TE = 2.45 ms; FOV = 256 × 232 mm; voxel resolution = 1.0 × 1.0 × 1.0-mm voxels. The rs-fMRI data were obtained with the following acquisition parameters: TR = 1000 ms; TE = 12, 29.75, 47.5 ms; multiband factor 4; flip angle = 50°; FOV = 240 × 240 mm; voxel resolution = 3.0 × 3.0 × 3.0-mm voxels.

### Preprocessing

Details of the HCP preprocessing can be found elsewhere (Van Essen et al., 2012a; Glasser et al., 2013; Smith et al., 2013). The HCP rs-fMRI data has been projected to fs_LR32k surface space (Van Essen et al., 2012b), denoised with ICA-FIX (Griffanti et al., 2014; Salimi-Khorshidi et al., 2014) and aligned with MSMAll (Robinson et al., 2014). Consistent with our previous studies (Li et al., 2019; He et al., 2020), we further applied global signal regression (GSR) and censoring to eliminate global and head motion related artifacts. More details of the processing can be found elsewhere (Li et al., 2019). After censoring, there might be less than 15 min of data for a particular run, but in the result section, we will still refer to uncensored frames as a 15-min run. We also used the ICA-FIX data in MNI152 space provided by the HCP. We then performed GSR, applied bandpass filtering (0.01 Hz – 0.1 Hz) and smoothed the data using a 4 mm full-width half maximum kernel in order to be consistent with previous work (Cash et al., 2021b). Runs with more than 50% censored frames were removed. Participants with all rs-fMRI runs remaining for all four sessions (N = 32) were considered.

In the case of the SING and SING-D datasets, we processed the data with the following steps. (1) Removal of the first 4 frames; (2) Slice time correction; (3) Motion correction and outlier detection: frames with FD > 0.2 mm or DVARS > 75 were flagged as censored frames. 1 frame before and 2 frames after these volumes were flagged as censored frames. Uncensored segments of data lasting fewer than five contiguous frames were also labeled as censored frames. Runs with over half of the frames censored were removed; After censoring, there might be less than 10 min of data for a particular run, but in the result section, we will still refer to uncensored frames as a 10-min run. (4) Correcting for susceptibility-induced spatial distortion; (5) Multi-echo denoising (DuPre et al., 2021); (6) Alignment with structural image using boundary-based registration (Greve and Fischl, 2009); (7) Global, white matter and ventricular signals, 6 motion parameters, and their temporal derivatives were regressed from the functional data. Regression coefficients were estimated from uncensored data; (8) Censored frames were interpolated with the Lomb-Scargle periodogram (Power et al., 2014); (9) Bandpass filtering (0.009 Hz – 0.08 Hz) was applied to the data; (10) The data was then projected onto FreeSurfer fsaverage6 surface space and smoothed using a 6 mm full-width half maximum kernel; (11) Lastly, preprocessed data in native fMRI volumetric space was smoothed using a 4mm full-width half maximum kernel. Alignment between T1 and MNI152 (FSL MNI152 2mm asymmetric template) was also computed using ANTs registration (Avants et al., 2011).

Because the HCP-processed data was provided in MNI152 space and also to be consistent with previous TMS targeting studies (Cash et al., 2021b), we opted to run all HCP analyses in MNI152 space. However, we note that in practice, personalized TMS is performed on the actual physical heads of participants, so personalized targets should ideally be estimated in native T1 space^1^. Therefore, in the case of the SING dataset, analyses were performed in individual native T1 space.

### Distance-to-scalp & sulcal depth maps

To favor targets closer to the scalp, we used distance-to-scalp as the primary anatomical metric. When direct distance-to-scalp measurements were not available, we substituted sulcal depth as a proxy for distance to the scalp. More specifically, to pick a target closer to the scalp, we computed the shortest distance from each gray matter voxel to the scalp in the SING dataset, yielding a distance-to-scalp brain map. A smaller distance indicates closer proximity to the scalp. In the case of the HCP dataset, recall our decision to perform the analysis in MNI152 space. Non-brain structures were removed in the HCP-provided data, so distance-to-scalp could not be computed. Instead, we used the sulcal depth map provided by FreeSurfer as a proxy for distance-to-scalp. During the preprocessing of the anatomical MRI data, FreeSurfer generates a highly accurate segmentation of the white matter and pial surfaces, as well as an estimate of the sulcal depth (average convexity) at each cortical location. A more negative sulcal depth (average convexity) value indicates closer proximity to the gyral crown, and thus presumably the scalp. We note that this assumption will be examined in a control analysis (see “Distance-to-scalp vs sulcal depth” in Methods).

### DLPFC mask & sACC time course for depression target localization

For depression target localization, we need to define a dorsolateral prefrontal cortex (DLPFC) mask and compute a sACC time course. The left DLPFC mask was defined as the union of spheres of 20-mm radii centered at BA9 (MNI x = -36, y = 39, z = 43), BA46 (MNI x = -44, y = 40, z = 29), the “5 cm rule” TMS site (MNI x = -41, y = 16, z = 54), and the Beam-F3 group-average stimulation site (MNI x = -37, y = 26, z = 49) (Fox et al., 2012; Cash et al., 2021b). Since the DLPFC mask was defined in MNI152 space, it can be directly used for the HCP dataset, whose analyses were performed in MNI152 space. To use the DLPFC mask in the SING dataset, the mask was projected from MNI space to the individual native T1 space for each participant.

To compute the sACC time series, the “group weight” method was used to alleviate dependency on the low signal-to-noise in the sACC region (Fox et al., 2013; Cash et al., 2021b). More specifically, 100 additional HCP participants underwent the same preprocessing procedure as the HCP test-retest dataset. We note that there was no overlap between the HCP test-retest participants and the 100 participants. The group-average sACC functional connectivity (FC) map was then computed based on the sACC ROI, which was defined as a sphere of 10-mm radius centered at MNI coordinates (x = 6, y = 16, z = -10) based on previous studies (Fox et al., 2012, 2013; Cash et al., 2021b).

For a new HCP participant in MNI space, the sACC ROI time course was defined as the weighted average of rs-fMRI time series, whose weights were given by the group-level sACC FC map. We note that only time courses of gray matter voxels (excluding the left DLPFC mask) contributed to this weighted average (Cash et al., 2021b). For a given SING participant, the procedure was the same except that the FC map was transformed from MNI space to individual native T1 space to compute the sACC ROI time course. The sACC ROI time course was then used to compute individual-level sACC FC map in either native T1 volumetric space (SING dataset) or MNI space (HCP dataset).

### Tree-based MS-HBM personalized depression target localization

Let us illustrate the tree-based MS-HBM algorithm using depression as an example. Furthermore, for the sake of this example, let us assume that we are working with a sulcal depth map like in the HCP dataset (Figures 1 and 2). We begin by applying MS-HBM to estimate individual-specific cortical networks from the preprocessed rs-fMRI data (Kong et al., 2019; Figure 1A). Since the dorsal attention and salience/ventral attention networks are known to be negatively correlated with the default network (Fox et al., 2006) and thus the sACC weighted time course (Fox et al., 2013), the portions of the individual-specific dorsal and salience/ventral attention networks within the DLPFC mask (from the previous section) are combined into a personalized MS-HBM ROI. Pseudocode for delineation of the personalized MS-HBM ROI is shown in Phase 1 of Figure 2.

**Figure 1.**
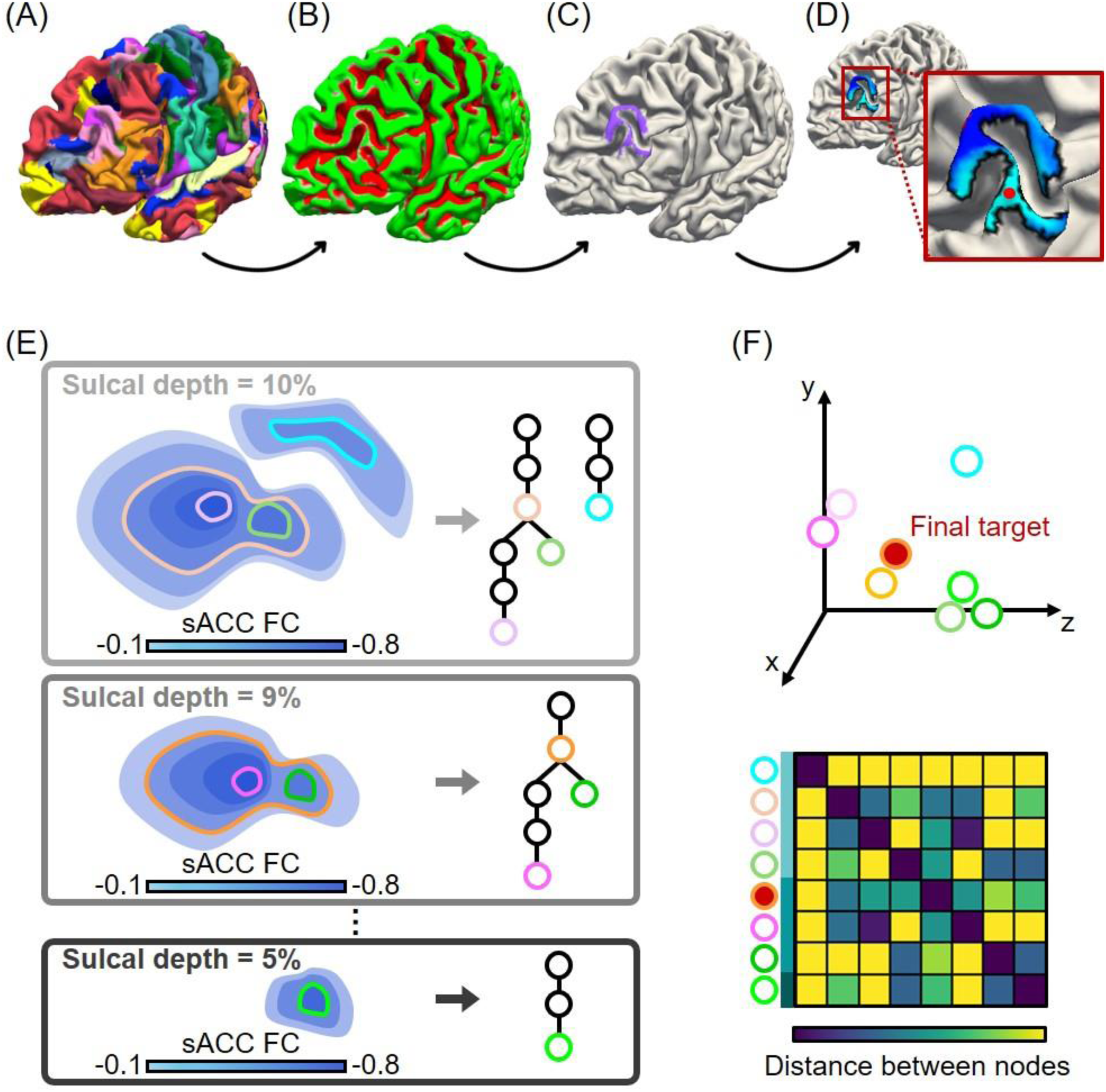
Illustration of tree-based MS-HBM personalized depression target localization in an example individual. (A) Individual-specific MS-HBM networks. (B) Individual-specific sulcal depth map. Green color indicates regions close to the gyral crowns (which are presumably close to the scalp). (C) The sulcal depth can be thresholded, retaining only the gyral portions of the individual’s salience/ventral attention and dorsal attention network components within DLPFC (shown in purple). (D) Color map shows FC (correlation) with sACC time course within the ROI from panel C, where blue indicates negative values. The FC map can be thresholded and the centroid of the largest connected component is used to obtain a personalized target. The difficulty of this simple approach is to choose an appropriate sulcal depth threshold (in panel C) and FC threshold (in panel D) that can generalize across individuals and datasets. Instead of using a fixed sulcal depth and FC thresholds, we propose a tree-based algorithm illustrated in (E) and (F). (E) Illustration of tree generation with different sulcal depth and sACC FC thresholds. White tree nodes with colored contours represent candidate targets. (F) A consensus target is obtained by finding the candidate that is closest in distance on average to all other candidate targets. Red node indicates the optimal target. Here we illustrate the tree-based algorithm with a sulcal depth map, but a distance-to-scalp map can also be used.

**Figure 2.**
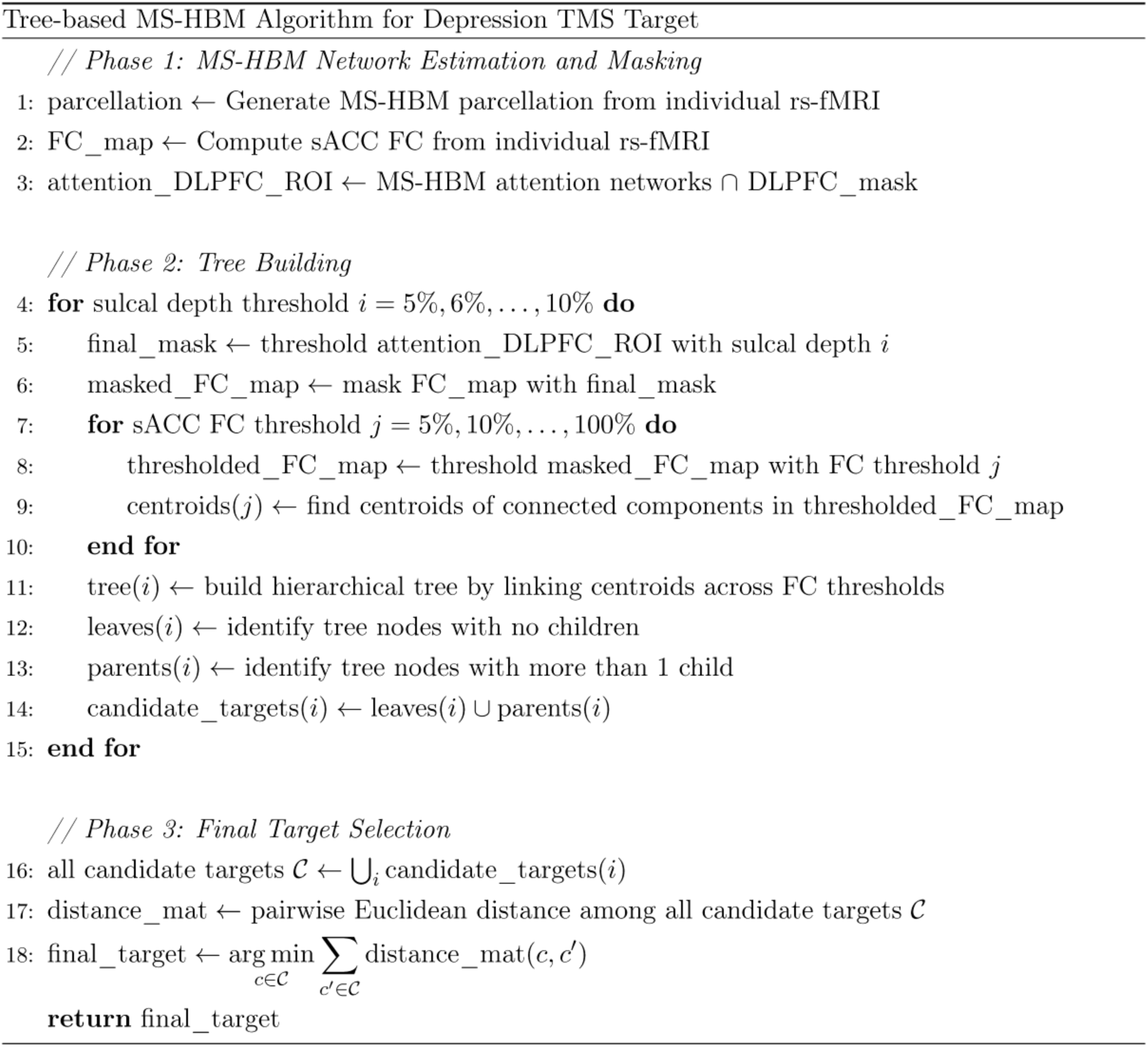
Pseudocode of tree-based MS-HBM for extracting a personalized depression target. The algorithm is divided into three phases. Functions and operations are described at a high level to illustrate the workflow and logic of the method.

To motivate our tree-based algorithm, let us first consider a simple approach of deriving a near-scalp depression target. Suppose we threshold the sulcal depth map (Figure 1B) within the personalized MS-HBM ROI, yielding a new personalized ROI (Figure 1C). The sACC FC map within this new ROI can then be computed (Figure 1D) and thresholded. The centroid of the largest component can be selected as the personalized target (Figure 1D). The drawback of this simple approach is that we have to choose (1) a threshold for the sulcal depth map, and (2) a threshold for the sACC FC map. Unfortunately, there is not an easy way to select these thresholds that are guaranteed to work well across datasets. For example, sACC FC values can vary significantly across datasets depending on the population, MRI acquisition and preprocessing procedures.

Instead, we propose a tree-based algorithm to address this issue. The idea is to define a reasonable range of values generalizable across a wide range of applications (e.g., datasets or disorders), and then find a consensus target location. More specifically, we define a range of sulcal depth values to be 5% to 10%. A x% threshold selects the top x% of brain locations closest to the gyral crown. We then vary the sulcal depth threshold from 5% to 10% in intervals of 1% (Figure 1E). We define the range of sACC correlation thresholds to be from 5% to 100%. A x% sACC threshold represents the top x% of brain locations most negatively correlated with sACC. We vary the sACC threshold from 5% to 100% in intervals of 5% (Figure 1E).

For a given sulcal depth threshold (e.g., 10%; top panel in Figure 1E) and the most stringent sACC threshold (i.e., 5%), connected components are extracted among brain voxels that survive the thresholds. The centroid of each connected component is then computed, yielding one or more centroids. These centroids form the leaves of a tree. As the sACC threshold becomes more lenient, the connected components grow bigger and new centroids are computed, which become the parents of the previous centroids. As the sACC threshold becomes more lenient, certain connected components from the previous threshold might merge. The centroid of the merged connected components becomes the parent centroid of the centroids from the previous connected components. Eventually, at the most lenient level, we may end up with one single connected component, whose centroid forms the root of the tree (middle panel of Figure 1E). However, multiple trees might also be obtained for a given sulcal depth threshold (top panel of Figure 1E).

Each sulcal depth threshold generates one or more trees (Figure 1E). For each tree, we select candidate targets corresponding to tree nodes with no children (i.e., leaves) and parent nodes with multiple children (colored nodes in Figure 1E). The leaves and parent nodes with multiple children are chosen as the candidate targets because they correspond to the centroids of connected components obtained from the strictest FC thresholds. Pseudocode for the generation of the candidate targets is shown in Phase 2 of Figure 2.

Finally, among all the candidates from all trees, a final target is then obtained by finding the candidate that is closest in distance on average to all other candidates (Figure 1F). Pseudocode for the final target generation is found in Phase 3 of Figure 2. The tree-based MS-HBM has essentially no parameter to tune. We do have to specify a range of sulcal depth (or distance-to-scalp) values and a range for FC values.

Throughout this study, we fixed the sulcal depth (or distance-to-scalp) range to be from 5% to 10% and the FC range to be from 5% to 100%. As a reminder, the target generation was performed in each individual’s native T1 volumetric space in the SING and SING-D datasets, and MNI volumetric space for the HCP dataset. However, for illustration, the procedure is shown on the cortical mid-thickness surface (Figure 1A-D). Finally, while the above procedure was illustrated with sulcal depth, we used distance-to-scalp measurements in the SING and SING-D datasets.

### Alternative approaches

We compared our tree-based approach with two approaches: the individualized cone algorithm (Fox et al., 2013), and the individualized connectome-guided cluster-based algorithm (Cash et al., 2021b).

The individualized cone algorithm extracted brain surface voxels 4 mm apart in the DLPFC mask (Fox et al., 2013). A series of concentric spheres of radius 2, 4, 7, 9, and 12 mm (referred to as “12 mm cone” in the original Fox study) centered on each voxel was chosen to match the linear decay of metabolic changes seen in animal TMS experiments (Valero-Cabré et al., 2005). The linear decay in the 12mm cone can be visualized in Supplementary Figure 2 of Fox and colleagues (2013). The varying intensity within the 12mm sphere was normalized to a mean of one and a weighted average was used to derive an overall sACC FC within this sphere. The optimal personalized location was the voxel location whose cone-weighted-average sACC FC was the most negative. By construction, the stimulation location should be close to the scalp.

The cluster algorithm used an individualized connectome-guided approach, reporting high reliability across sessions (Cash et al., 2021b). Briefly, the cluster algorithm thresholded the sACC FC map within a DLFPC mask. The centroid of the largest component was then selected as the personalized target. However, the cluster approach does not restrict the target to be near the scalp, so the resulting targets might be deep in the sulcus.

### Quantitative evaluation of depression targets: single fMRI run

We applied tree-based MS-HBM, cluster and cone algorithms to extract depression targets independently from the first session and third session of the SING dataset. As a reminder, we note that the first and third sessions were two weeks apart. Recall that each session contains a single 10-min rs-fMRI run. The procedure was repeated independently for the first run of the first session and the first run of the third session in the HCP dataset. As a reminder, we note that the first and third sessions were on average 11.6 months apart. Recall that each HCP run was 14 min 33 sec long. We considered three evaluation metrics.

The first metric we considered was distance-to-scalp or sulcal depth. We computed distance between target and scalp in the SING dataset. T1 from the third session was used to compute distance-to-scalp for the target derived from the first session. The procedure was repeated by reversing the sessions and averaging the evaluation metric. Since distance-to-scalp was not available in the HCP dataset, we computed sulcal depth instead. Once again, T1 data from the third session was used to compute sulcal depth for the target derived from the first session, and vice versa. The procedure was repeated by reversing the sessions and averaging the evaluation metrics. A smaller distance-to-scalp (or more negative sulcal depth) indicates closer proximity to the scalp (or gyral crown), and thus better performance.

The second metric was inter-individual/intra-individual distance, which is a measure of reliability (Cash et al., 2021b). More specifically, intra-individual distance was simply defined as the Euclidean distance between the individual’s targets derived from the first MRI session and the third MRI session. A lower intra-individual distance indicates better reliability, but it is not a sufficiently good metric by itself. The reason is that a group-level target has an intra-individual distance of zero, suggesting perfect intra-individual agreement. However, a group target would also have no inter-individual variability (see Figure S1 for illustration). Therefore, we also computed inter-individual distance for each individual, which was defined as the average Euclidean distance between the individual’s personalized target and other individuals’ personalized targets. In the case of the SING dataset, the inter-individual distance was computed for each individual by transforming other individuals’ targets into the native T1 volumetric space of the individual. In the case of the HCP datasets, no transformation was necessary since all targets were already in MNI152 space. Finally, inter-individual/intra-individual distance was defined as the ratio of inter-individual distance and intra-individual distance. A larger inter-individual/intra-individual distance indicates greater reliability, and thus better performance.

The third metric is inter-session sACC FC (Fox et al., 2013; Cash et al., 2021b). More specifically, for a given individual, the target from the first session was used as a seed region to compute the FC with the sACC time course in the third session. The procedure was repeated by reversing the sessions and averaging the evaluation metrics. A more negative sACC FC indicates better performance.

We note that the cluster algorithm required the specification of a sACC FC threshold. Here, we performed a leave-one-individual-out cross-validation within each dataset to determine the optimal threshold for each individual. As an example, to derive the personalized cluster target of the first HCP individual, we tested a whole range of FC thresholds for the remaining 31 HCP participants to optimize the sACC FC. The FC threshold that yielded the best (most negative) sACC FC was used to derive the personalized target of the first HCP individual. This procedure was repeated for each individual.

### Impact of rs-fMRI scan duration

Longer scan duration improves the quality of brain parcellations (Laumann et al., 2015; Gordon et al., 2017; Kong et al., 2019, 2021), as well as individual-level prediction of phenotypic traits (Feng et al., 2023; Ooi et al., 2025). Here, we repeated the analyses from the previous section by varying the number of rs-fMRI runs.

In the previous section, we used the first and third sessions in the SING dataset, i.e., 10 min of fMRI data was used to derive personalized targets independently in each session. Here we repeated the analyses using the first two sessions and the last two sessions, i.e., 20 min of fMRI data was used to derive personalized targets independently in the first two sessions and last two sessions.

In the HCP dataset, we previously performed the analyses using the first run of the first session and first run of the third session, i.e., 14 min 33 sec of fMRI data was used to derive personalized targets independently each session. Here we increased the number of runs, so we ended up using ∼15 min, ∼29 min, ∼44 min or ∼58 min of data to derive personalized targets.

### Distance-to-scalp vs sulcal depth

As previously mentioned, the HCP analyses were performed using sulcal depth instead of distance-to-scalp measurements because non-brain structures were already removed from the HCP-provided data. On the other hand, the analyses in the SING dataset utilized distance-to-scalp measurements. As a control analysis, we examined whether sulcal depth was a good proxy of distance-to-scalp by repeating tree-based MS-HBM in the SING dataset using sulcal depth and a 10-min rs-fMRI run. We then re-examined the evaluation metrics: (1) distance-to-scalp, (2) inter/intra-distance reliability, and (3) sACC FC.

### Control analysis with the dysphoric target map

Instead of using the sACC weight map, we repeated the previous analysis using the dysphoric target map (Siddiqi et al., 2020) in the SING dataset. One key difference is that instead of minimizing the target’s FC with the weighted sACC time course (i.e., we wanted the FC to be more negative), here we maximized the target’s FC with the weighted dysphoric target map time course (i.e., we wanted the FC to be more positive).

### Tree-based MS-HBM personalized anxiosomatic target localization

To demonstrate the versatility of the tree-based MS-HBM algorithm, we applied the same algorithm without any parameter tuning to extract personalized anxiosomatic targets in the SING dataset. Instead of the previously defined DLPFC mask for depression targets, we defined a dorsal PFC (DPFC) mask by creating a 30-mm radius sphere around the group-level anxiosomatic target (MNI x = 18, y = 37, z = 55) and intersecting that with the MNI template. Instead of the sACC weight map, we used the anxiosomatic circuit map (Siddiqi et al., 2020; Taylor et al., 2024). However, instead of minimizing the target’s FC with sACC (i.e., we wanted the FC to be more negative), here we seek to maximize the target’s FC with the anxiosomatic time course computed from the anxiosomatic circuit map (i.e., we wanted the FC to be more positive). Finally, because the anxiosomatic circuit map overlapped with control-B, default-A and default-B group-level networks from our previous study (Kong et al., 2019), the personalized MS-HBM ROI comprised the individual-specific control-B, default-A and default-B networks within the new DPFC mask. All tree-based MS-HBM parameters remained the same between the depression and the anxiosomatic target localization.

### Quantitative evaluation of targets in patients with treatment-resistant depression

We applied tree-based MS-HBM, cluster and cone algorithms to extract depression targets independently from the first run and second run of the SING-D dataset. Both runs were collected in the same session and lasted 10 minutes each. Consistent with the analyses in the SING and HCP datasets, we computed the distance-to-scalp metric, inter-individual/intra-individual distance reliability metric, and the inter-session sACC FC metric for SING-D participants.

### E-field analyses in patients with treatment-resistant depression

We also compared the electric field (E-field) properties of tree-based MS-HBM targets with cluster and cone targets in the SING-D dataset. Individualized head models were generated from the anatomical MRI scans using SimNIBS 4.1.0 (Thielscher et al., 2015). For each participant, we simulated the E-field distribution for both test and retest targets using the MagVenture Cool-B65 coil model with coil angle of 45 degree. To assess target quality, three evaluation metrics were considered.

For the first two evaluation metrics, the “E-field hotspot” was defined as the set of cortical vertices representing the top X percent of E-field magnitude (Lynch et al., 2022; Ren et al., 2025), with X ranging from 0.1 to 1. For each participant and each targeting approach, one run was used to generate the E-field hotspot based on the target estimated from that run. The other run – which we will refer to as the “unseen” run – was then used to compute the evaluation metrics. The roles of the two runs were then swapped and the metrics were then averaged across the two runs.

The first evaluation metric was “network specificity”, which was defined as the percentage of the E-field hotspot area occupied by the target networks (dorsal and salience/ventral attention networks) of the unseen run (Lynch et al., 2022; Ren et al., 2025). Second, we computed the FC between the E-field hotspot and the sACC using data from the unseen run (Fox et al., 2013; Cash et al., 2021b; Elbau et al., 2023). More specifically, a rs-fMRI time course was computed based on a weighted average of E-field hotspot vertices’ time courses from the unseen run, where the weights corresponded to the local E-field strength. The hotspot time course was then correlated with the sACC weighted time course from the same run. A more negative hotspot sACC FC might indicate better target quality (Weigand et al., 2018; Elbau et al., 2023).

Finally, the third evaluation metric was the reliability of the induced E-fields by computing inter-individual/intra-individual E-field correlations in the SING-D dataset. More specifically, intra-individual E-field correlation was simply defined as the correlation between the E-field maps of individual’s targets derived from the first run and the second run. A higher intra-individual correlation indicates better reproducibility. To compute the inter-individual E-field correlation for a particular individual, we first transformed other individuals’ targets into the native T1 volumetric space of the individual, and then generated the E-field maps of the transformed targets. Correlations between the individual’s E-field map and the E-field maps of the transformed targets were then computed and averaged. A lower inter-individual correlation indicates higher inter-individual variability. Finally, inter-individual/intra-individual correlation was defined as the ratio of inter-individual correlation and intra-individual correlation. A smaller inter-individual/intra-individual correlation indicates greater E-field reliability, and thus better performance.

### Data and code availability

This raw data for HCP test-retest dataset is publicly available (https://www.humanconnectome.org/). Code for this work is freely available at the GitHub repository maintained by the Computational Brain Imaging Group (https://github.com/ThomasYeoLab/CBIG). Processing pipelines of the fMRI data can be found here (https://github.com/ThomasYeoLab/CBIG/tree/master/stable_projects/preprocessing/CBIG_fMR <u>I_Preproc2016</u>). The individual-specific MS-HBM parcellation approach can be found here (https://github.com/ThomasYeoLab/CBIG/tree/master/stable_projects/brain_parcellation/Kong2 <u>019_MSHBM</u>). Code specific to the analyses in this study can be found here (https://github.com/ThomasYeoLab/Kong2026_TMSTree).

## Results

### Reliable near-scalp personalized depression targets derived from tree-based MS-HBM

Tree-based MS-HBM was used to derive a personalized depression target in each SING participant’s native T1 volumetric space using a 10 min rs-fMRI scan from the first MRI session. The personalized targets from the 18 SING participants were then projected to the fsaverage6 surface space to visualize individual differences in target locations (Figure 3A). Most targets were located in the group-level salience / ventral attention networks within DLPFC (Kong et al., 2019). However, some targets were in the group-average control-A and control-C network, suggesting that the individual-specific attention networks of these participants extended beyond the group-level attentional network boundaries.

**Figure 3.**
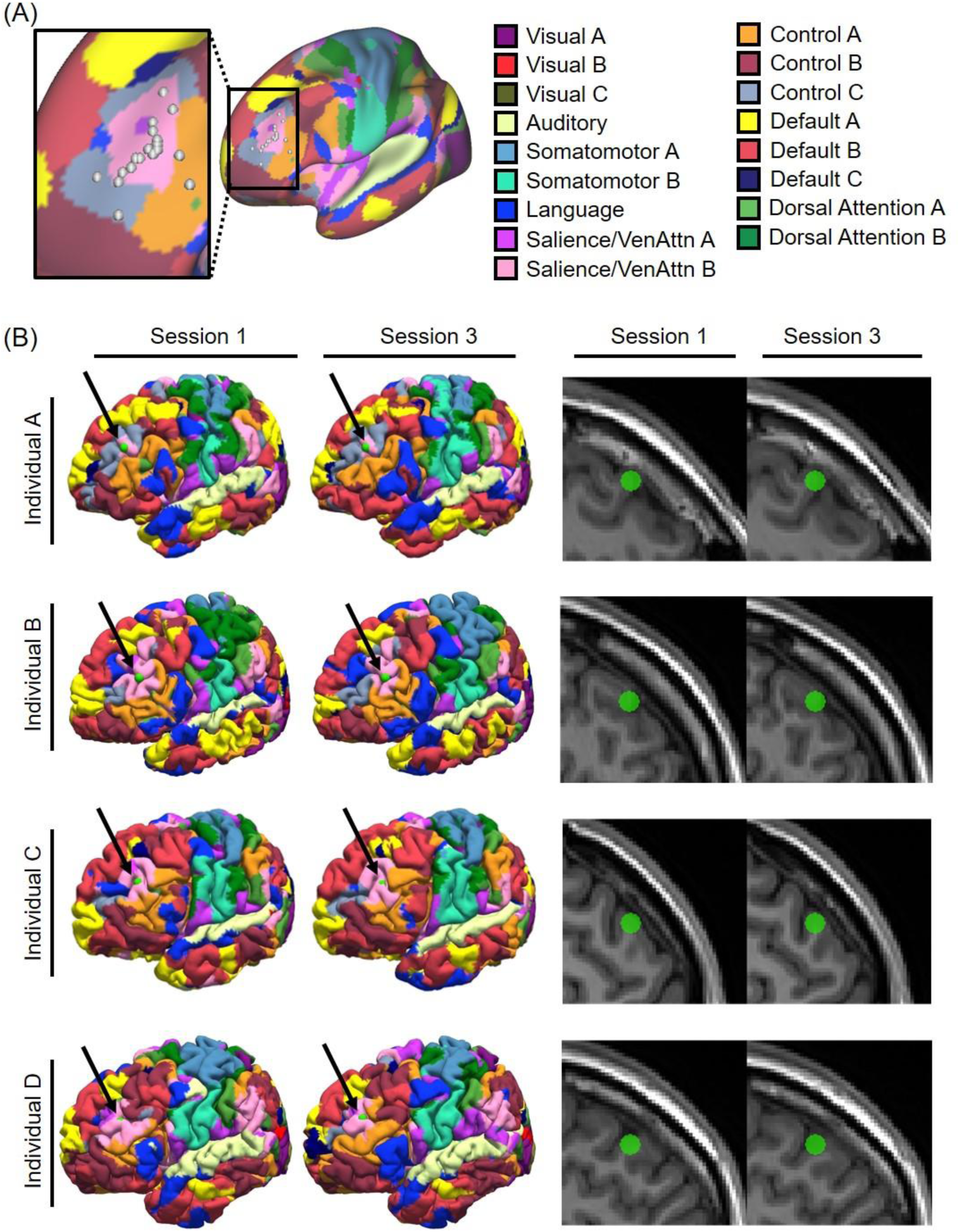
Reliability of personalized depression targets derived from tree-based MS-HBM. (A) Personalized depression targets of 18 SING participants overlaid on a group-level map of 17 resting-state networks (Kong et al., 2019). Target locations were estimated in each participant’s native T1 volumetric space using the 10 min rs-fMRI scan from the first MRI session and then projected to fsaverage6 surface space for visualization. (B) Tree-based MS-HBM targets are near the scalp and demonstrate high within-individual reliability across two sessions (two weeks apart) in the SING dataset. Four representative participants are shown here, with targets (green circles) displayed on native pial surfaces (left) and native T1 images (right). Black arrows point to the green circles on the native pial surfaces. A target located deep within a sulcus would not be visible on the native pial surface, so the visible targets (green circles) indicate that tree-based MS-HBM was able to successfully localize near-scalp targets.

We also estimated targets using the 10 min rs-fMRI scan from the third session (2 weeks after the first session). Visual inspection suggests that the MS-HBM networks exhibited significant inter-individual variability, while being reliable within each individual across sessions (Figure 3B left panel). Within-individual variability in target locations across the two sessions was on average 4.3 mm (as measured in native volumetric space).

The target locations were all close to the scalp (Figure 3B right panel) with an average distance of 13.9 mm from the scalp as measured in native volumetric space. Participant A provides an illustrative example where the individual-specific attention networks were buried deep in the sulci in DLPFC, with only a small portion of the networks extending onto the gyri (Figure 3A left panel). Tree-based MS-HBM was able to reliably identify a personalized target within the attention networks that was close to the scalp in both sessions.

### Tree-based MS-HBM depression targets achieved more negative sACC FC than alternative approaches

Here we compared tree-based MS-HBM, cluster (Cash et al., 2021b), and cone (Fox et al., 2013) targets in terms of closeness to the scalp (or gyral crown), test-retest reliability, and sACC FC strength. More specifically, all three algorithms were used to derive depression targets independently from the first session and third session of the SING dataset. Each SING session contained a single 10-min rs-fMRI run. The procedure was repeated independently for the first run of the first session and the first run of the third session in the HCP dataset. Each HCP run was 14 min 33 sec long.

We first evaluated the pairwise Euclidean distances between targets derived from the different targeting algorithms. In the HCP dataset, the distances were on average 11.3 ± 6.5 mm (mean ± std) between tree-based MS-HBM and cluster targets, 11.3 ± 10.4 mm between tree-based MS-HBM and cone targets, and 11.1 ± 6.1 mm between cone and cluster targets (Figure S2A). In the SING dataset, the corresponding distances were 13.1 ± 7.8 mm, 12.8 ± 10.0 mm, and 12.5 ± 9.9 mm, respectively (Figure S2B).

Not surprisingly, in the HCP dataset, tree-based MS-HBM and cone targets were closer to the gyral crowns (and presumably the scalp) than the cluster targets (p < 3.1e-5; Figure 4A). Tree-based MS-HBM and cone targets were also closer to the scalp than the cluster targets in the SING dataset (p < 6.9e-5; Figure 4B). We remind the reader that targets were computed in one session and distance-to-scalp (or sulcal depth) was computed in the other session. The procedure was repeated by reversing the sessions and averaging the evaluation metrics.

**Figure 4.**
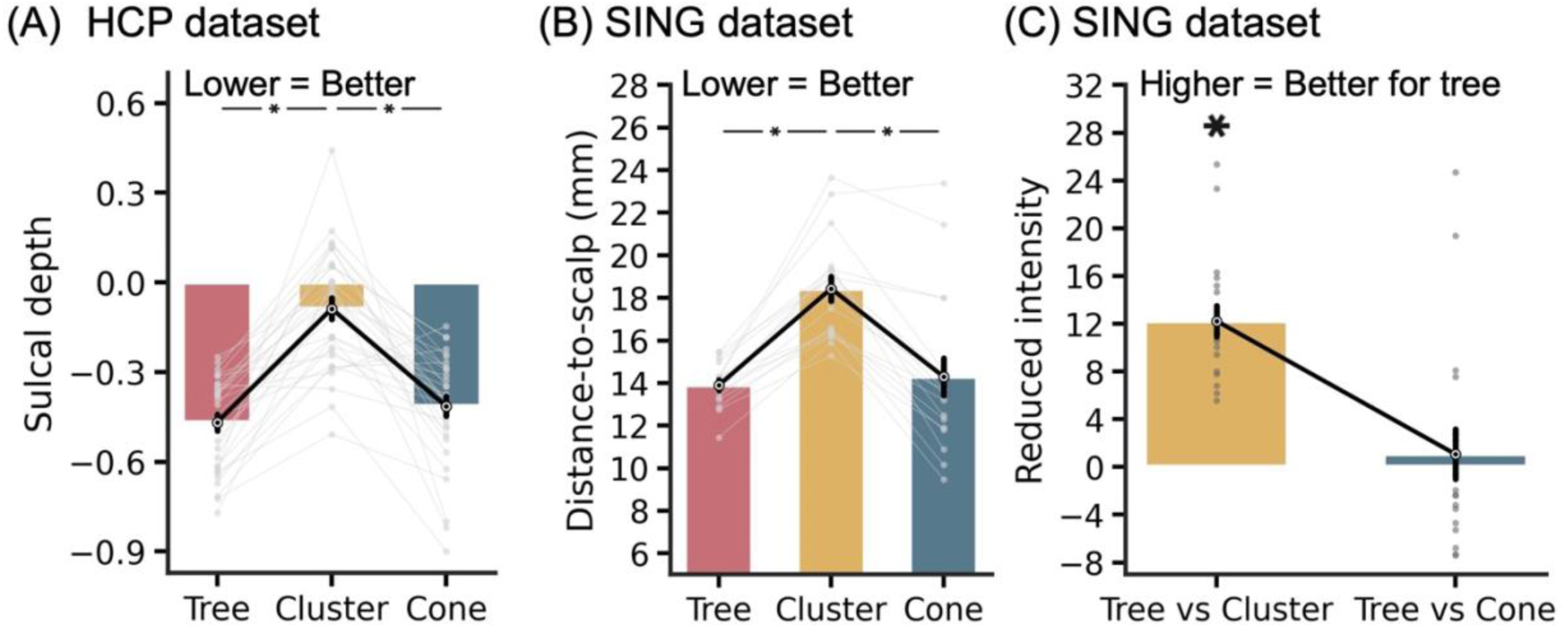
Comparison of tree-based MS-HBM, cluster and cone targets in terms of closeness to the scalp (or gyral crown) and TMS intensity. (A) Sulcal depth (average convexity) in the HCP dataset. We remind the reader that non-brain structures were removed from the data provided by the HCP, so distance-to-scalp could not be computed. Instead, sulcal depth (average convexity) was used as a proxy. A more negative value indicates closer distance to the gyral crown and presumably the scalp (see Figure 7). (B) Distance-to-scalp in the SING dataset. A smaller value indicates better performance. (C) Reduction in stimulation intensity in the SING dataset. Following the SNT protocol (Cole et al., 2020), by using a linear adjustment in stimulation intensity based on distance-to-scalpand assuming 90% RMT dosage, a hypothetical reduction in stimulation intensity between tree-based MS-HBM and other approaches can be computed. Lower intensity indicates better performance. * indicates statistical significance after multiple comparisons correction with FDR q < 0.05. We note that in all analyses, targets were derived in one session and then evaluated in another session.

In the case of the SING dataset, following the SNT protocol (Cole et al., 2020), a linear adjustment was used to convert distance-to-scalp measurements into TMS intensity, assuming 90% RMT dosage. Based on the relationship reported by Stokes and colleagues, a one-mm reduction in distance-to-scalp corresponds to a 3% decrease in TMS intensity at 100% RMT, which scales to 2.7% reduction per millimeter at 90% RMT^2^ (Stokes et al., 2005). The tree-based MS-HBM targets led to significantly lower stimulation intensity than the cluster targets, with an average reduction of 12.2% (Figure 4C).

In terms of reliability, tree-based MS-HBM exhibited numerically better (greater) inter-individual/intra-individual distance than the cluster algorithm with statistical significance achieved in the HCP dataset (p = 0.034; Figure 5A) but not in the SING dataset (Figure 5B). Tree-based MS-HBM and cone exhibited similar reliability.

**Figure 5.**
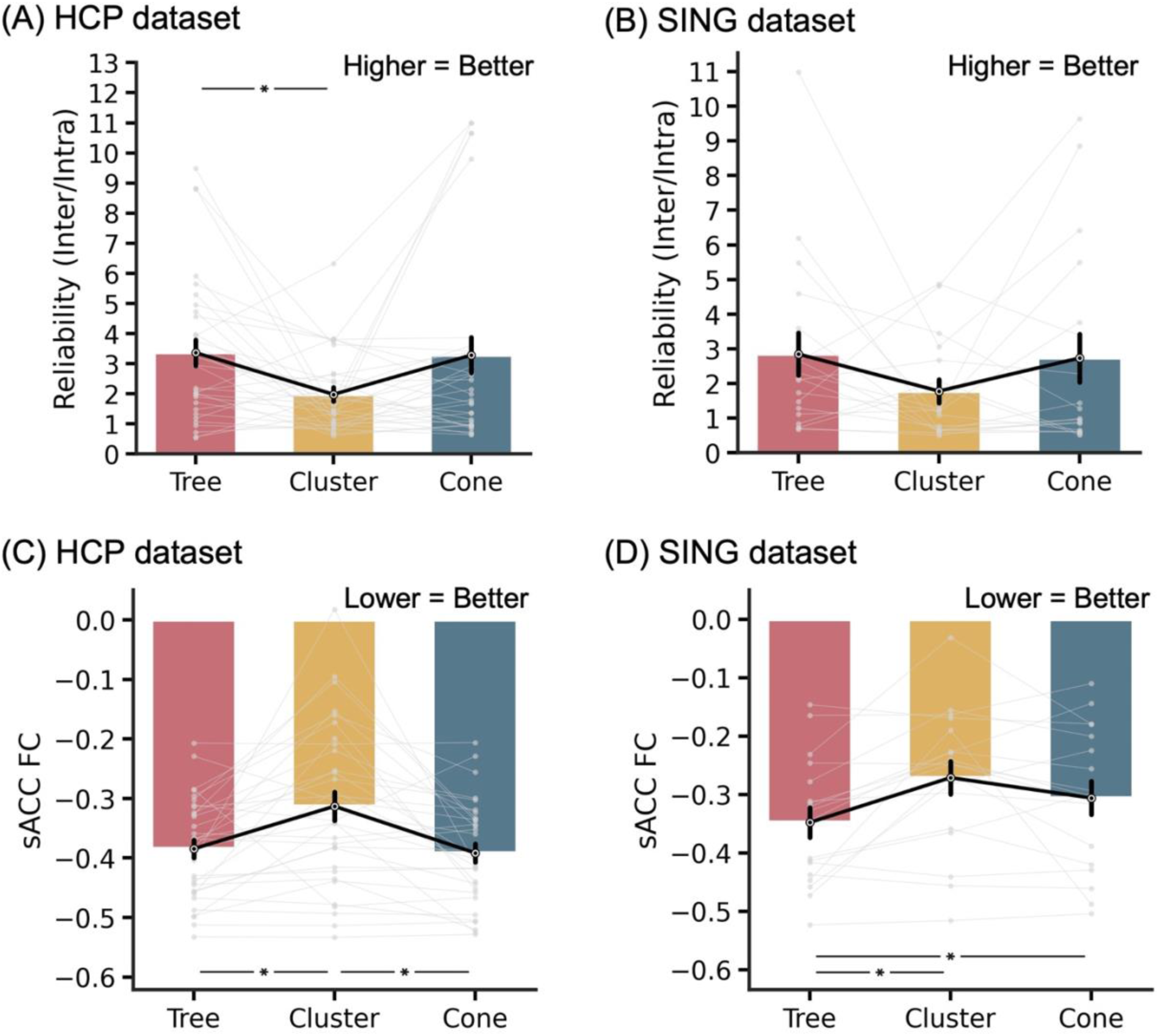
Comparison of tree-based MS-HBM, cluster and cone targets in terms of test-retest reliability and sACC FC in two independent datasets. (A) Reliability (ratio of inter-individual distance and intra-individual distance) in the HCP dataset. A higher value indicates better performance. (B) Reliability (ratio of inter-individual distance and intra-individual distance) in the SING dataset. (C) sACC FC in the HCP dataset. A more negative value indicates better performance. (D) sACC FC in the SING dataset. * indicates statistical significance after multiple comparisons correction with FDR q < 0.05. We note that in all analyses, targets were derived in one session and then evaluated in another session.

In the case of sACC FC (Figures 5C & 5D), tree-based MS-HBM targets exhibited statistically better (more negative) sACC FC than the cone targets in the SING dataset (p = 0.036), but not the HCP dataset (p = 0.514). The MS-HBM also exhibited statistically better sACC FC than the cluster algorithm in both SING (p = 0.018) and HCP (p = 0.018) datasets. As a reminder, targets were computed in one session and sACC FC was evaluated in the other session. The procedure was repeated by reversing the sessions and averaging the evaluation metrics.

### Regardless of scan duration, tree-based MS-HBM depression targets compared favorably with alternative approaches

To explore the effect of scan duration, we repeated the analyses from the previous section using more rs-fMRI runs (Figure 6). Not surprisingly, regardless of scan duration, tree-based MS-HBM and cone targets were closer to the gyral crowns (and presumably the scalp) than the cluster targets in the HCP dataset (p < 4.2e-5; Figure 6A). Tree-based MS-HBM and cone targets were also closer to the scalp than the cluster targets in the SING dataset (p < 6.9e-5; Figure 6B).

**Figure 6.**
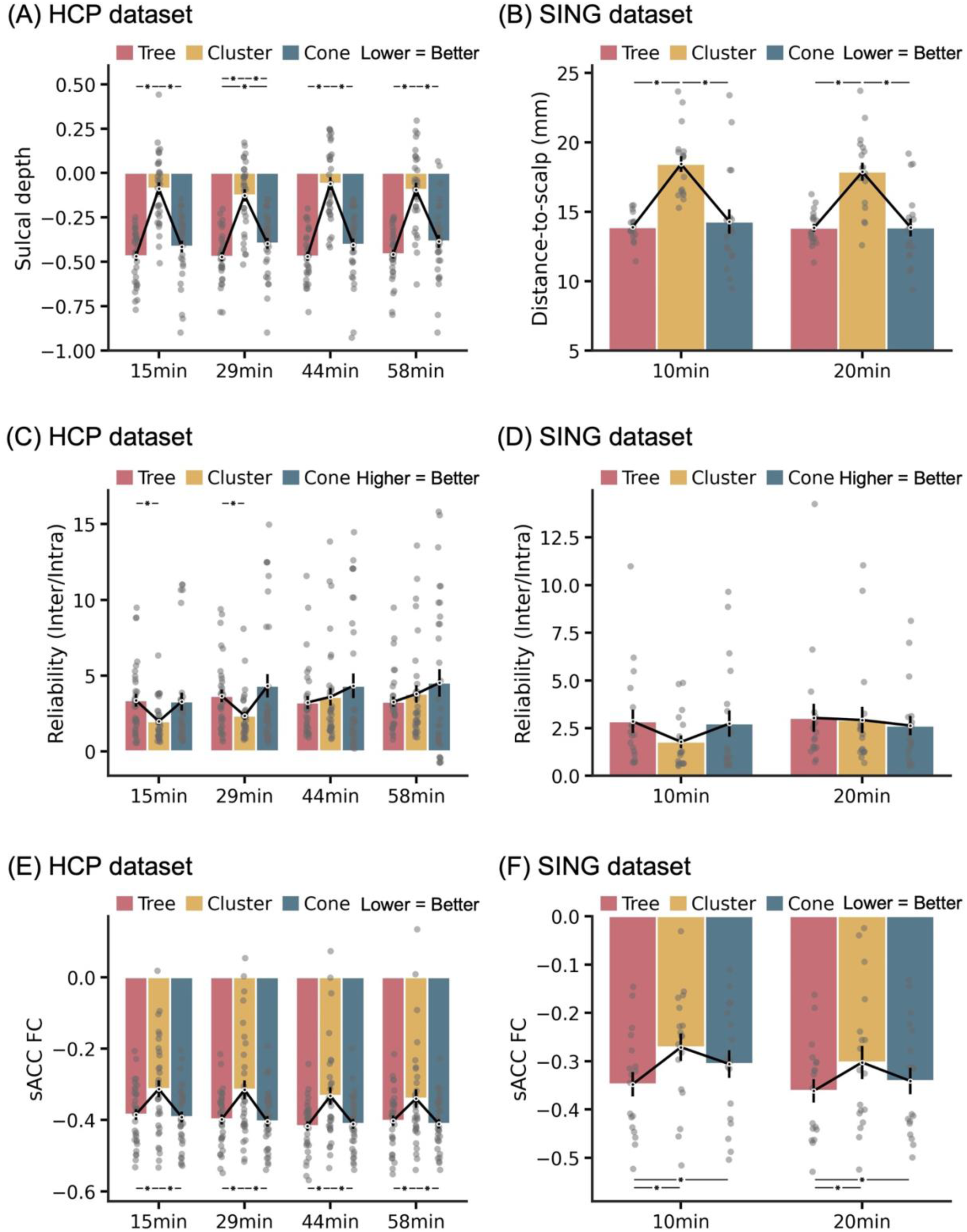
Impact of rs-fMRI scan duration on evaluation metrics. (A) Sulcal depth in the HCP dataset. A more negative value indicates closer distance to the gyral crown and presumably the scalp (see Figure 6). (B) Distance-to-scalp in the SING dataset. A smaller value indicates better performance. (C) Reliability (ratio of inter-individual distance and intra-individual distance) in the HCP dataset. A higher value indicates better performance. (D) Reliability (ratio of inter-individual distance and intra-individual distance) in the SING dataset. (E) sACC FC in the HCP dataset. A more negative value indicates better performance. (F) sACC FC in the SING dataset. * indicates statistical significance after multiple comparisons correction with FDR q < 0.05. We note that in all analyses, evaluation was performed on leave-out sessions that were not used to derive the targets.

Tree-based MS-HBM targets were also more reliable than the cluster targets for one rs-fMRI run (15 min; p = 0.034) and two rs-fMRI runs (29 min; p = 0.012) in the HCP dataset. With three or more rs-fMRI runs (44 min or longer), the cluster algorithm exhibited numerically better reliability than tree-based MS-HBM, but the improvement was not significant (Figure 6C). In the SING dataset, both tree-based MS-HBM and cluster targets exhibited similar reliability with 20 min of rs-fMRI (Figure 6D).

There was no statistical difference in reliability between tree-based MS-HBM and cone targets in both HCP and SING datasets (Figures 6C and 6D). Similar to cluster targets, the cone targets exhibited higher reliability with longer scan duration in the HCP dataset (Figure 6C). However, although cone targets exhibited numerically better reliability than tree-based MS-HBM for two or more rs-fMRI runs (29 min or longer), the improvement was not significant (Figure 6C). In the SING dataset, both tree-based MS-HBM and cone targets exhibited similar reliability with 20 min of rs-fMRI.

Finally, in the case of sACC FC, tree-based MS-HBM targets were statistically better than the cluster targets even up to 58 min of rs-fMRI (Figures 5E & 5F). There was no statistical difference in sACC FC between tree-based MS-HBM and cone targets in the HCP dataset across all scan durations (Figure 6E). In the SING dataset, tree-based MS-HBM targets exhibited statistically better sACC FC than cone targets across all scan durations (Figure 6F).

### Sulcal depth is a reasonably good proxy of distance-to-scalp

The HCP analyses were performed using sulcal depth instead of distance-to-scalp measurements because distance-to-scalp measurements were unavailable. As a control analysis, the tree-based MS-HBM algorithm was applied to derive depression targets independently from the first session and third session of the SING dataset, using either distance-to-scalp or sulcal depth maps. As can be seen for a representative individual (Figure 7A), the sulcal depth and distance-to-scalp maps are highly correlated. Indeed, the reliability (inter/intra-individual distance), sACC FC and distance-to-scalp were also similar between distance-to-scalp and sulcal-depth targets (Figures 7B to 7D). Therefore, sulcal depth map is a good proxy for distance-to-scalp measurements when the latter are unavailable (like in the HCP dataset). However, in practice, personalized targets should be derived in native volumetric space, so there is no reason not to use actual distance-to-scalp measurements.

**Figure 7.**
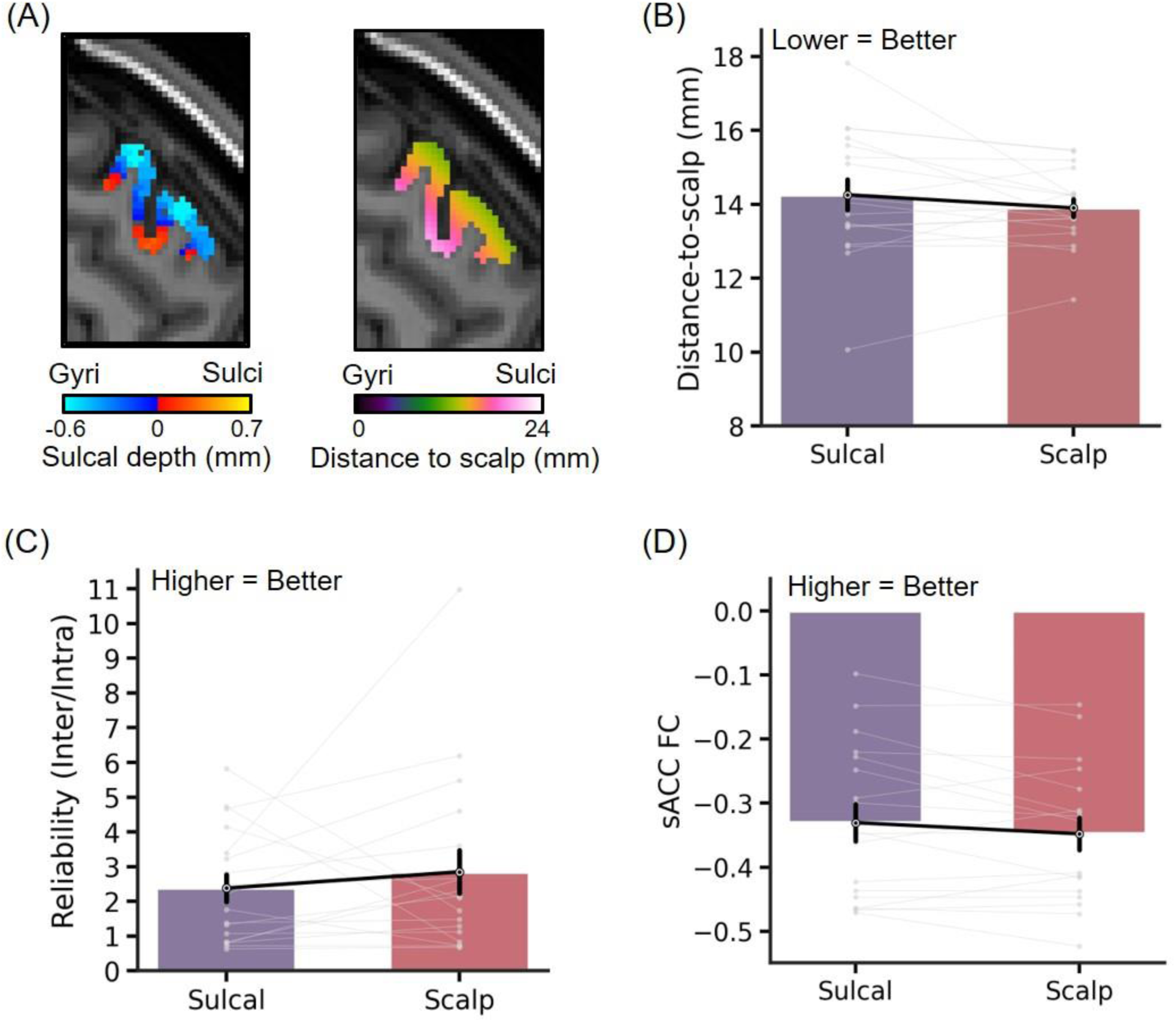
Comparison of evaluation metrics when sulcal depth or distance-to-scalp maps were used to generate tree-based MS-HBM depression targets in the SING dataset. (A) Sulcal depth map and distance-to-scalp map within the personalized MS-HBM ROI in an example individual. (B) Comparison of target distance-to-scalp measurements (C) Comparison of reliability (ratio of inter-individual distance and intra-individual distance). A higher value indicates better performance. (D) Comparison of sACC FC. A more negative value indicates better performance. There was no statistical significance after multiple comparisons correction with FDR q < 0.05.

### Control analysis with the dysphoric target map

Instead of the sACC weight map, we considered the use of the dysphoric target map (Siddiqi et al., 2020) in the SING dataset. Similar to Figures 4 and 5, targets were estimated in the first session and evaluated in the third session (and vice versa).

Consistent with previous results (Figure 4), both tree-based MS-HBM and cone targets were closer to the scalp than the cluster algorithm (Figure S3B). Based on the SNT protocol (Stokes et al., 2005; Cole et al., 2020), this translated to 14.6% reduction in stimulation intensity for the tree-based MS-HBM targets compared with the cluster targets (Figure S3C).

Consistent with previous results (Figure 5), Tree and cone algorithms exhibited numerical better reliability than the cluster algorithms, but the differences were not statistically significant (Figure S3D). Finally, tree-based MS-HBM targets exhibited statistically better (higher) dysphoric FC than both cluster and cone targets (Figure S3E).

### Personalized anxiosomatic targets derived from tree-based MS-HBM

To demonstrate the versatility of the tree-based MS-HBM algorithm, we applied the same algorithm, without tuning any parameter, to extract personalized anxiosomatic targets in the SING dataset. The group-level anxiosomatic circuit map was derived from a previous study (Siddiqi et al., 2020; Figure 8A). Based on a single rs-fMRI run (10-min), we observed significant inter-individual variability in personalized tree-based MS-HBM targets when visualized on the fsaverage6 surface space (Figure 8B).

**Figure 8.**
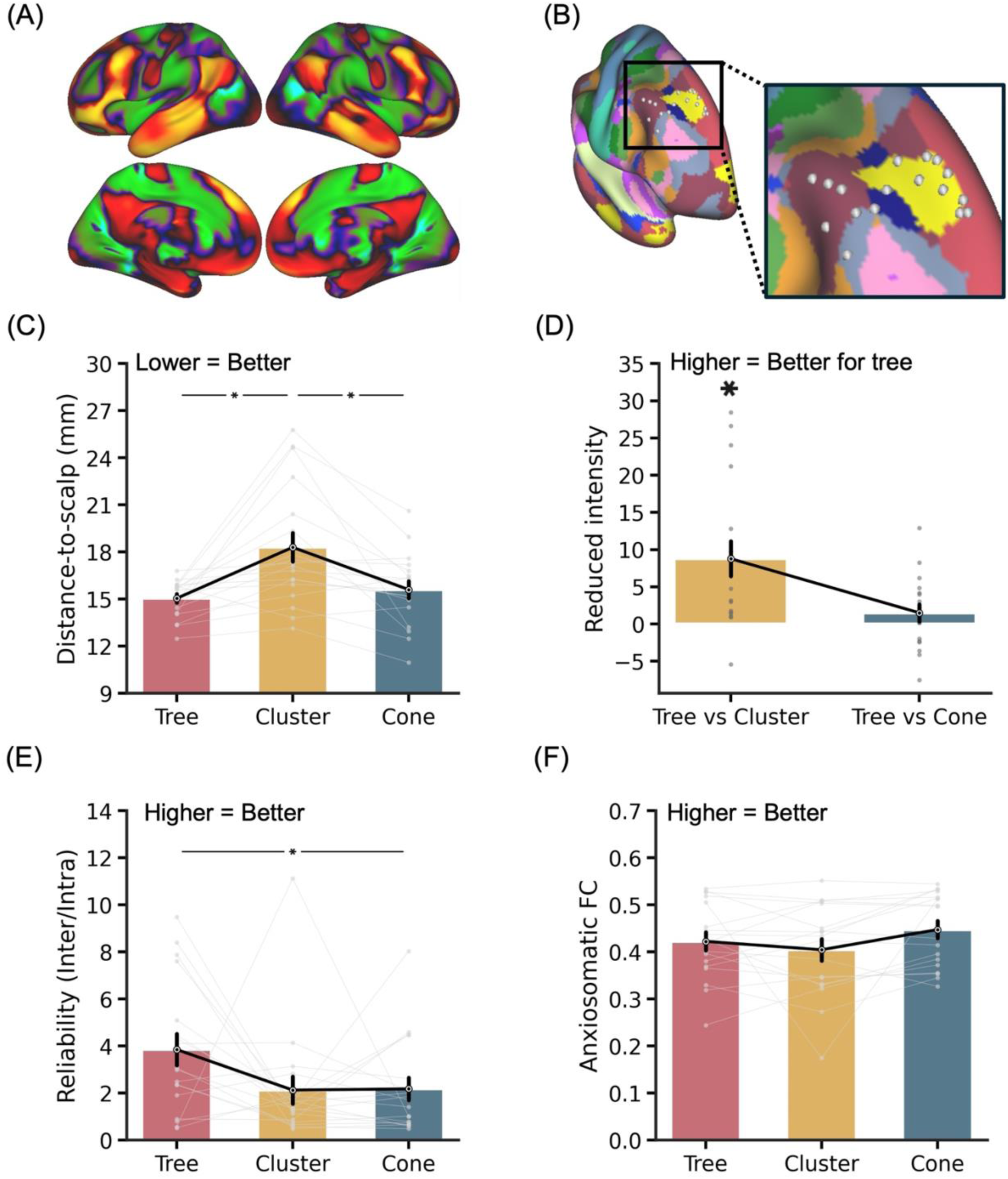
Comparison of personalized anxiosomatic targets generated by tree-based MS-HBM, cluster and cone algorithms in the SING dataset. (A) Group-level anxiosomatic circuit map visualized on fsaverage6 surface. (B) Personalized anxiosomatic targets of 18 SING participants overlaid on a group-level map of 17 resting-state networks (Kong et al., 2019). Target locations were estimated in each participant’s native T1 volumetric space using the 10 min rs-fMRI scan from the first MRI session and then projected to fsaverage6 surface space for visualization. (C) Distance-to-scalp in the SING dataset. A smaller value indicates better performance. (D) Reduction in stimulation intensity in the SING dataset. Following the SNT protocol (Cole et al., 2020), by using a linear adjustment in stimulation intensity based on distance-to-scalp and assuming 90% RMT dosage, a hypothetical reduction in stimulation intensity between tree-based MS-HBM and other approaches can be computed. (E) Reliability (ratio of inter-individual distance and intra-individual distance) in the SING dataset. A higher value indicates better performance. (F) FC with the anxiosomatic circuit map in the SING dataset. A more positive value indicates better performance. * indicates statistical significance after multiple comparisons correction with FDR q < 0.05. We note that in all analyses, targets were derived in one session and then evaluated in another session.

Tree-based MS-HBM targets were significantly closer to the scalp than the cluster targets (p = 0.017; Figure 8C), which translated to 8.8% lower TMS intensity (Figure 8D). While the tree-based MS-HBM targets exhibited greater reliability than the cluster targets, the difference was not significant (Figure 8E). On the other hand, tree-based MS-HBM targets were statistically more reliable than the cone targets (p = 0.039; Figure 8E). Across the three approaches (tree-based MS-HBM, cluster and cone), cone targets exhibited the best (most positive) FC with the anxiosomatic circuit map (Figure 8F), but all differences were not significant.

### Tree-based MS-HBM outperforms cluster and cone in treatment-resistant depression

To evaluate whether the tree-based MS-HBM algorithm generalized to a clinical population, we applied the tree-based MS-HBM, cone, and cluster algorithms to the SING-D dataset comprising 43 patients with treatment-resistant depression. The tree-based MS-HBM and cluster targets were on average 12.7 ± 6.2 mm (mean ± std) apart. The tree-based MS-HBM and cone targets were on average 12.9 ± 13.1 mm apart, while cone and cluster targets were on average 13.8 ± 9.0 mm apart (Figure S2C).

The tree-based MS-HBM targets were significantly closer to the scalp than the cone and cluster targets (p < 4e-4; Figure 9A), corresponding to 15.4% lower TMS intensity relative to cluster targets and 4.5% lower intensity relative to cone targets based on the SNT protocol (Figure 9B). Tree-based MS-HBM also demonstrated the highest reliability, achieving statistical significance against the cluster targets (p = 0.037), but not the cone targets. In the case of sACC FC, tree-based MS-HBM targets exhibited statistically better (more negative) sACC FC than both cone and cluster targets in the SING-D dataset (p < 0.025; Figure 9D).

**Figure 9.**
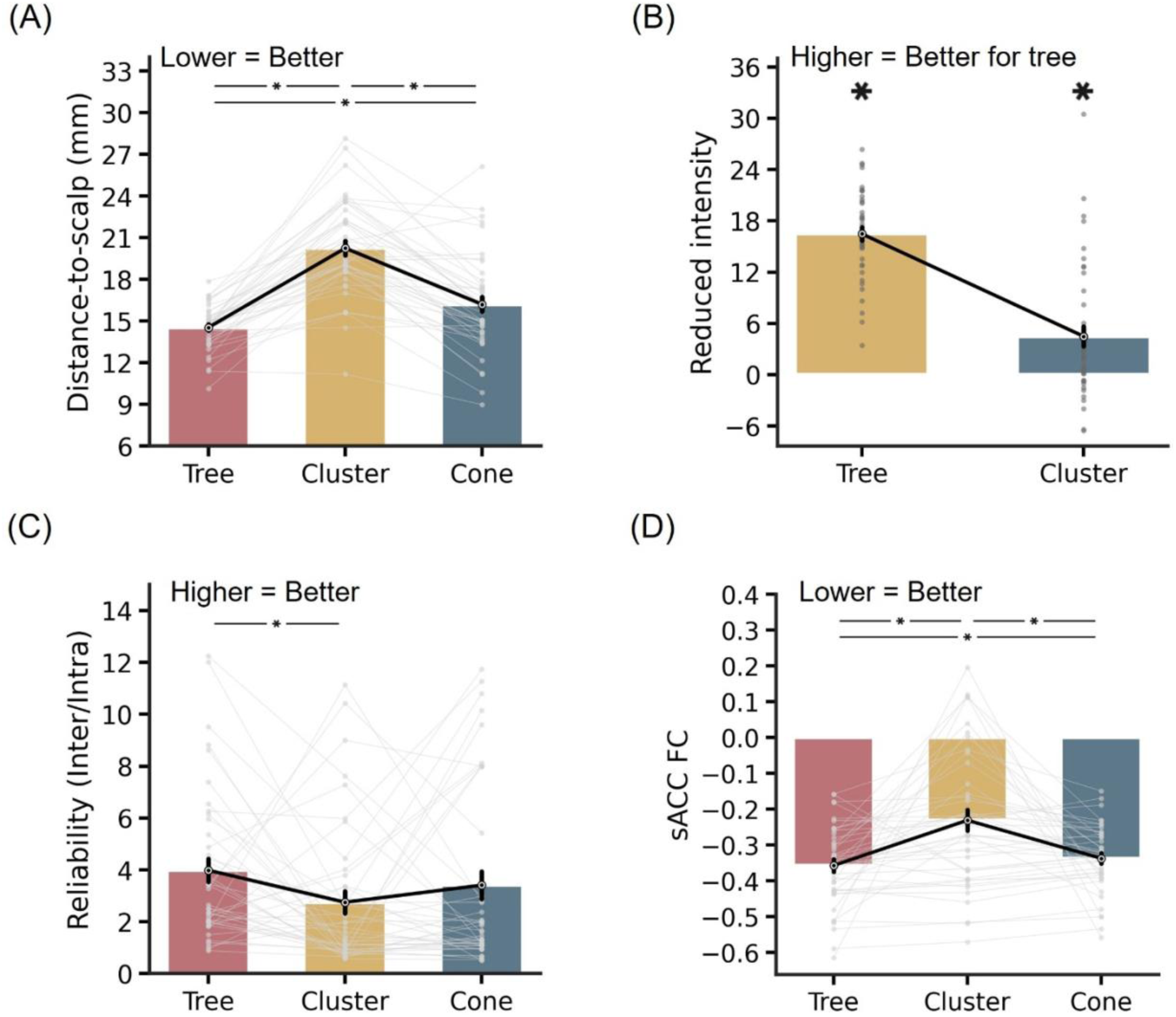
Comparison of personalized depression targets generated by tree-based MS-HBM, cluster and cone algorithms in the SING-D dataset, comprising 43 patients with treatment-resistant depression. (A) Distance-to-scalp in the SING-D dataset. A smaller value indicates better performance. (B) Reduction in stimulation intensity in the SING-D dataset. Following the SNT protocol (Cole et al., 2020), by using a linear adjustment in stimulation intensity based on distance-to-scalp and assuming 90% RMT dosage, a hypothetical reduction in stimulation intensity between tree-based MS-HBM and other approaches was computed. (C) Reliability (ratio of inter-individual distance and intra-individual distance) in the SING-D dataset. A higher value indicates better performance. (D) sACC FC in the SING-D dataset. A more negative value indicates better performance. * indicates statistical significance after multiple comparisons correction with FDR q < 0.05. We note that in all analyses, targets were derived in one run and then evaluated in another run.

Figure 10A illustrates the E-field characteristics of the tree-based MS-HBM target of a representative individual from the SING-D dataset. By construction, tree-based MS-HBM algorithm identifies a personalized target within attentional networks in left DLPFC that exhibited the strongest negative FC with sACC. Therefore, unsurprisingly, tree-based MS-HBM targets produced significantly stronger on-target E-field engagement of attentional networks in the unseen run compared with both cone and cluster targets across different E-field hotspot thresholds (p < 0.018; Figure 10B).

**Figure 10.**
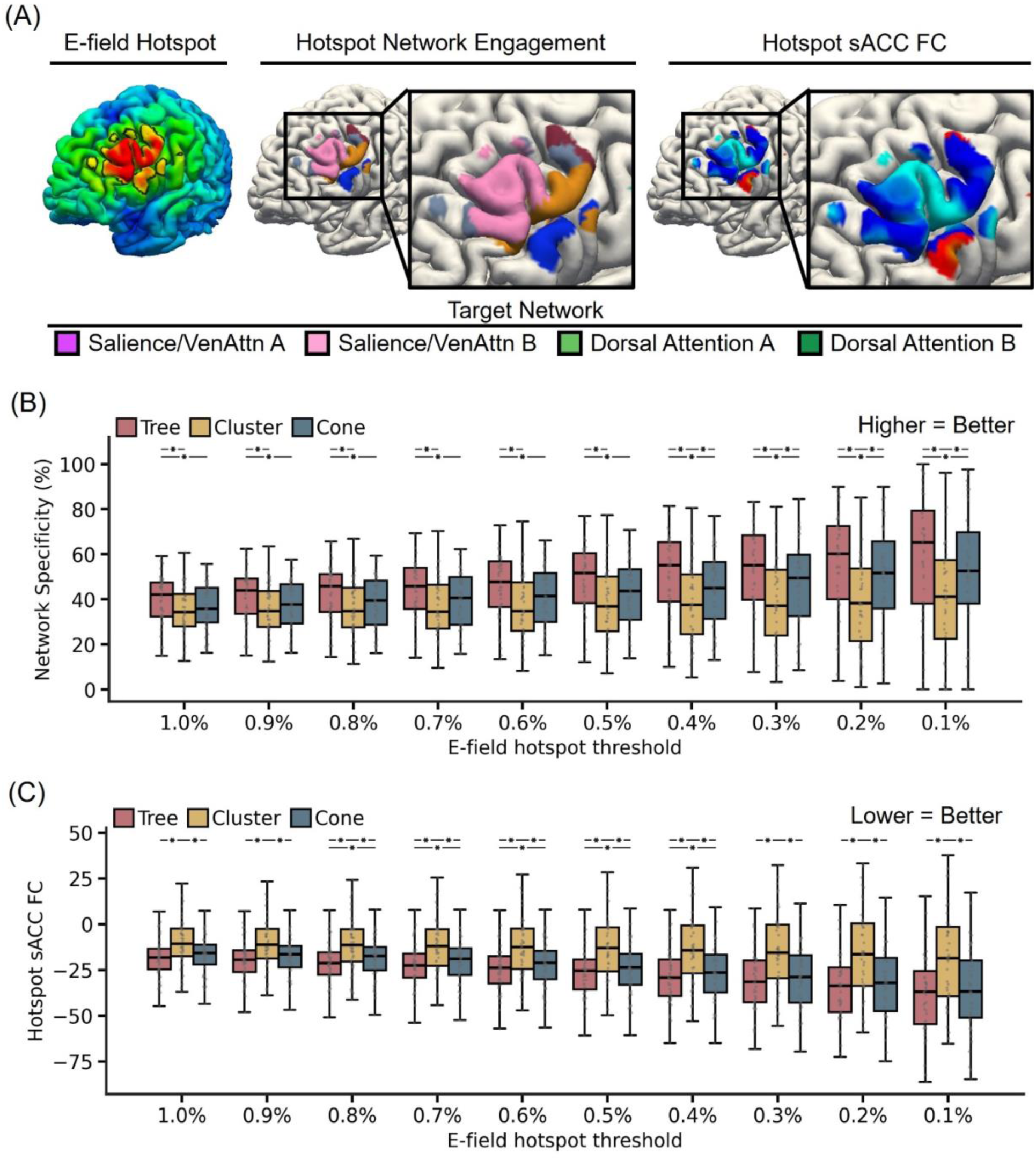
Comparison of network specificity and hotspot sACC FC based on electric field modeling of tree-based MS-HBM targets, cone, and cluster targets in the SING-D dataset, comprising 43 patients with treatment-resistant depression. (A) E-field characteristics of the tree-based MS-HBM target of a representative participant. The left panel shows the E-field maps estimated using SimNIBS, with the “E-field hotspot” defined as the top 1 percent of cortical vertices (black boundaries) following previous studies. The middle panel shows the personalized MS-HBM networks within the E-field hotspot The right panel shows functional connectivity of sACC within the E-field hotspot. (B) Comparison of tree-based MS-HBM, cluster, and cone targets in terms of attentional network E-field engagement at E-field hotspot thresholds ranging from 1% to 0.1%. Here attention networks referred to salience/ventral attention networks and dorsal attention networks. Higher values indicate greater engagement of the attention networks. (C) Comparison of tree-based MS-HBM, cluster, and cone targets in terms of hotspot sACC FC at E-field hotspot thresholds ranging from 1% to 0.1%. Lower values indicate stronger anticorrelation with sACC. * indicates statistical significance after multiple comparisons correction with FDR q < 0.05. We note that in all analyses, targets were derived in one run and then evaluated in another run.

In addition, the E-field hotspot of tree-based MS-HBM targets also exhibited stronger (more negative) hotspot sACC FC in the unseen run than both cone and cluster targets, with statistical significance observed across all E-field hotspot thresholds for cluster targets (p < 9e-5) and across E-field hotspot thresholds from 0.4% to 0.8% for cone targets (p < 0.033; Figure 10C). These results suggest that tree-based MS-HBM targets might more strongly modulate the sACC circuit.

Figure 11A shows the E-field maps from both runs of two representative participants for tree-based MS-HBM, cluster, and cone targets. Tree-based MS-HBM targets exhibited significantly higher intra-individual E-field similarity than both cluster and cone targets (p < 2.3e-3; Figure 11B), indicating improved within-subject reproducibility. Inter-individual E-field variability did not differ across algorithms (Figure 11C). Importantly, the inter-individual/intra-individual E-field correlation ratios of tree-based MS-HBM targets were significantly higher than both cone and cluster targets (p < 0.031; Figure 11D), suggesting better reliability of the induced E-fields.

**Figure 11.**
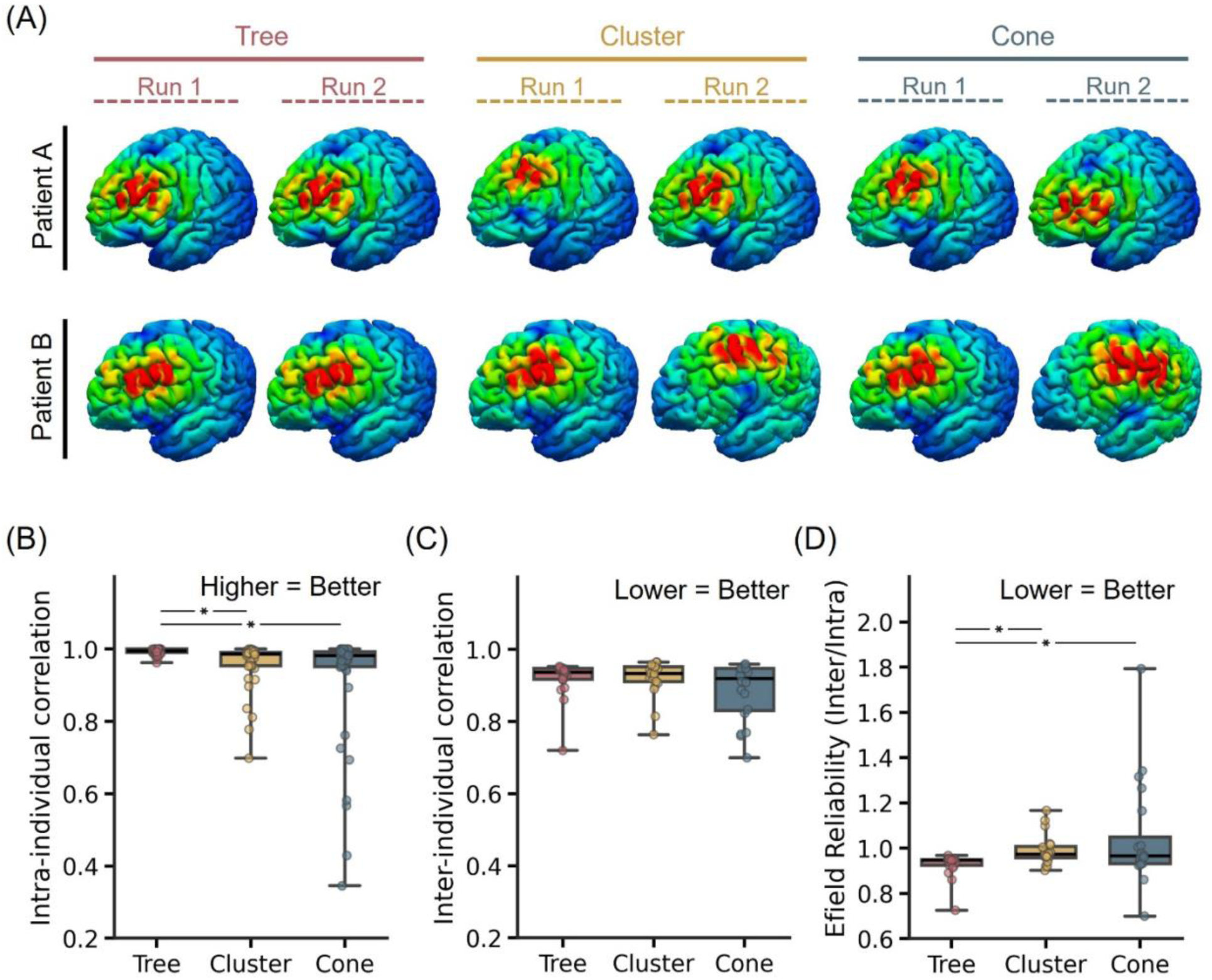
E-field reliability of personalized depression targets generated by tree-based MS-HBM, cone, and cluster algorithms in the SING-D dataset, comprising 43 patients with treatment-resistant depression. (A) E-field maps of targets from two different runs of two representative SING-D participants for tree-based MS-HBM, cluster, and cone targets. (B) Intra-individual E-field correlation across runs. A higher value indicates greater within-individual similarity. (C) Inter-individual E-field correlation across participants. A lower value indicates higher variability across participants. (D) E-field reliability (ratio of inter-individual correlation and intra-individual correlation). A lower value indicates better performance.

## Discussion

In this study, we proposed and evaluated the tree-based MS-HBM algorithm for deriving personalized brain stimulation targets. A key advantage of the tree-based algorithm is the elimination of pre-defined FC or scalp-proximity thresholds. Therefore, the resulting tree-based MS-HBM algorithm has essentially no “free” parameter to tune.

Compared to the cluster algorithm (Cash et al., 2021b), tree-based MS-HBM depression targets were significantly closer to the scalp, yielding 12.2% lower stimulation intensity (based on a hypothetical SNT protocol). Tree-based MS-HBM targets were generally more reliable than the cluster targets, while exhibiting better (more negative) sACC FC. Tree-based MS-HBM’s superior scalp proximity and sACC FC persisted even with longer fMRI scan durations. On the other hand, longer scan durations led to comparable reliability between tree-based MS-HBM and cluster targets. In the case of the anxiosomatic target, both algorithms have similar reliability and anxiosomatic FC, but the tree-based MS-HBM targets were again closer to the scalp, yielding 8.6% lower stimulation intensity (based on a hypothetical SNT protocol).

On the other hand, tree-based MS-HBM and cone (Fox et al., 2013) depression targets exhibit similar proximity to scalp and reliability. However, tree-based MS-HBM targets exhibited better (more negative) sACC FC. The advantage in sACC FC persisted even with longer scan durations. In the case of the anxiosomatic target, both targets were similarly close to the scalp. Cone targets exhibited numerically better FC, but the difference was not significant. On the other hand, the tree-based MS-HBM targets were statistically more reliable than the cone targets.

In a dataset of individuals with treatment-resistant depression, tree-based MS-HBM depression targets were significantly closer to the scalp than cluster and cone targets, yielding 15% and 5% lower stimulation intensity respectively (based on a hypothetical SNT protocol). Tree-based MS-HBM targets also exhibited the best (most negative) sACC FC. Finally, tree-based MS-HBM targets also exhibited the most specific engagement of attentional networks, best (most negative) E-field hotspot sACC FC, and induced the most reliable E-field. Therefore, tree-based MS-HBM targets were not only closer to the scalp but also produced E-field characteristics that better align with hypothesized therapeutic circuits, supporting their potential physiological relevance.

### Scan duration & MS-HBM

Early rs-fMRI studies focused on group-level networks (Damoiseaux et al., 2006; Power et al., 2011; Yeo et al., 2011). However, advances in MRI acquisition (Setsompop et al., 2012; Lynch et al., 2020) and dense-sampling studies (Laumann et al., 2015; Braga and Buckner, 2017; Gordon et al., 2017) reveal that every individual exhibits their own unique network topography. Individual network topography corresponds well to individual-specific task activation patterns (Seitzman et al., 2019; Braga et al., 2020; Du et al., 2024), predicts cognitive, personality and mental health traits (Bijsterbosch et al., 2018; Kong et al., 2019), and is more similar in monozygotic twins than dizygotic twins (Anderson et al., 2021).

However, long rs-fMRI scans are clinically impractical due to cost and patient compliance. Algorithmic advances (Harrison et al., 2015; Kong et al., 2019) now enable high-quality individual-specific network estimation from shorter scans. Our MS-HBM explicitly differentiates within-individual from between-individual variability. By ignoring within-individual variability, previous network mappings might confuse within-individual variability with between-individual variability.

With just 10-min scans, MS-HBM is able to estimate networks of equivalent quality as other approaches using 50 mins of fMRI (Kong et al., 2019). MS-HBM parcellations generalize better to *new* rs-fMRI and task-fMRI from the same individuals than other algorithms (Kong et al., 2019; Kong et al., 2021). The strong generalization property of MS-HBM might be why tree-based MS-HBM targets exhibited better sACC FC in a new fMRI session, compared with cluster and cone targets.

### Choice of MS-HBM networks for personalized target selection

At the group-level, the salience/ventral attention networks (but not the dorsal attention networks) are present in the DLPFC (Figure 3; Yeo et al., 2011; Kong et al., 2019). Notably, the salience network is enlarged in depression (Lynch et al., 2024). While this might suggest defining a personalized ROI based solely on the salience/ventral attention networks, both dorsal attention and salience/ventral attention networks are strongly anticorrelated with the default network (Fox et al., 2005, 2006), and thus sACC. Furthermore, in some individuals, small portions of the dorsal attention networks do appear in DLPFC. Therefore, we defined the personalized ROI based on both dorsal attention and salience/ventral attention networks.

On the other hand, the group-level anxiosomatic circuit map appeared similar to the default network (Siddiqi et al., 2020). However, on closer inspection, we found that the anxiosomatic circuit map overlapped with control-B and default networks in our 17-network group-level atlas from our previous study (Kong et al., 2019). Consequently, in the current study, the personalized ROI included individual-specific control-B and default networks. It is possible that better treatment outcomes might be achieved by constraining the personalized ROI solely to the default networks. Future studies – whether via retrospective TMS analyses (Cash et al., 2021a; Siddiqi et al., 2021) or prospective trials – will be necessary to determine the optimal networks for defining the personalized ROI.

### Limitations & future work

Scalp distance and inter-individual/intra-individual reliability were utilized as surrogate validation metrics rather than predictors of antidepressant efficacy. These metrics reflected practical and regulatory considerations relevant to the clinical translation of TMS targeting strategies. Scalp distance influences stimulation intensity and therefore impacts patient tolerability and treatment retention. Reliability is a fundamental requirement for analytical validation in regulatory frameworks (e.g., FDA guidance on software as a medical device). Algorithms that produce highly variable targets across sessions are unlikely to be considered generalizable or clinically deployable, regardless of efficacy observed in a single dataset.

Negative sACC FC was utilized as a hypothesized circuit-level marker motivated by prior literature, although it has not been conclusively established as a biomarker of antidepressant response. However, we emphasize that the proposed tree-based MS-HBM framework is biomarker-agnostic: its primary contribution is to provide a principled and reliable method for translating any hypothesized circuit map into a personalized, near-scalp TMS target. If future work demonstrates that alternative biomarkers – such as positive sACC FC (Oathes et al., 2023) – are superior, the same framework can be readily applied. Thus, while the optimal biomarker for depression remains an active area of investigation, tree-based MS-HBM is designed to be robust to evolving biological hypotheses and adaptable to future discoveries.

Future work may leverage retrospective analyses of patients who previously received anatomically guided TMS (e.g., Cash et al., 2021a; Siddiqi et al., 2021) to determine whether stimulation locations that lie closer to tree-based MS-HBM targets are associated with superior clinical outcomes relative to cluster-based or cone-based targeting approaches. In parallel, we have completed an open-label clinical study evaluating tree-based MS-HBM targets for depression (Kong et al., 2026) and are currently conducting a double-blind trial directly comparing tree-based MS-HBM targeting with the Beam-F3 approach (NCT06385223). Additional prospective studies could extend this framework by contrasting personalized anxiety targets with group-level anxiety maps or Beam-F3–based targeting.

Finally, in our current framework, distance-to-scalp was used as a practical constraint to favor targets that are more accessible to stimulation, rather than a biological surrogate for efficacy. Our post-hoc evaluations (Figures 9 and 10) confirmed that these clinically accessible targets yield robust E-field engagement of the intended therapeutic circuits, supporting their potential physiological relevance. However, it is important to note that geometric proximity alone might not guarantee effective simulation. E-field modeling may offer superior precision for individualized dosing (Numssen et al., 2024) and target selection (Weis et al., 2020; Balderston et al., 2022; Lynch et al., 2022; Lueckel et al., 2023; Dannhauer et al., 2024; Sun et al., 2024). Integrating such modeling directly into the tree-based MS-HBM framework is a high-priority avenue for future iterations, potentially further enhancing circuit-based neuromodulation.

## Conclusion

We introduced the tree-based MS-HBM algorithm for deriving personalized near-scalp, connectome-guided brain stimulation targets. Overall, when considering all three metrics (reliability, FC and scalp proximity), tree-based MS-HBM was comparable or better than both cluster (Cash et al., 2021) and cone (Fox et al., 2013) algorithms for delineating depression and anxiosomatic targets. These results demonstrate the versatility of the tree-based MS-HBM framework for deriving personalized stimulation targets across psychiatric conditions.

## Author Contributions

RK: Conceptualization, Methodology, Software, Validation, Project administration, Writing - Original Draft. AX: Methodology, Software. LQRO: Software, Validation. CA, XWT, SEG, JJL, JZJK, RSYT, HKGS: Data acquiring. TWKT: Software. RDW: Validation. MDF, SS: Conceptualization, Methodology. PCT: Conceptualization, Methodology, Data acquiring, Funding acquisition. BTTY: Conceptualization, Methodology, Supervision, Funding acquisition. All authors revised the paper and provided critical feedback.

## Acknowledgement

Our research is supported by the NUS Yong Loo Lin School of Medicine (NUHSRO/2020/124/TMR/LOA), the Singapore National Medical Research Council (NMRC) LCG (OFLCG19May-0035), NMRC CTG-IIT (CTGIIT23jan-0001), NMRC OF-IRG (OFIRG24jan-0006; OFIRG24jul-0049), NMRC STaR (STaR20nov-0003), Singapore Ministry of Health (MOH) Centre Grant (CG21APR1009), the Temasek Foundation (TF2223-IMH-01), and the United States National Institutes of Health (R01MH133334). Any opinions, findings and conclusions or recommendations expressed in this material are those of the authors and do not reflect the views of the funders.

## Supplementary Material

**Figure S1.**
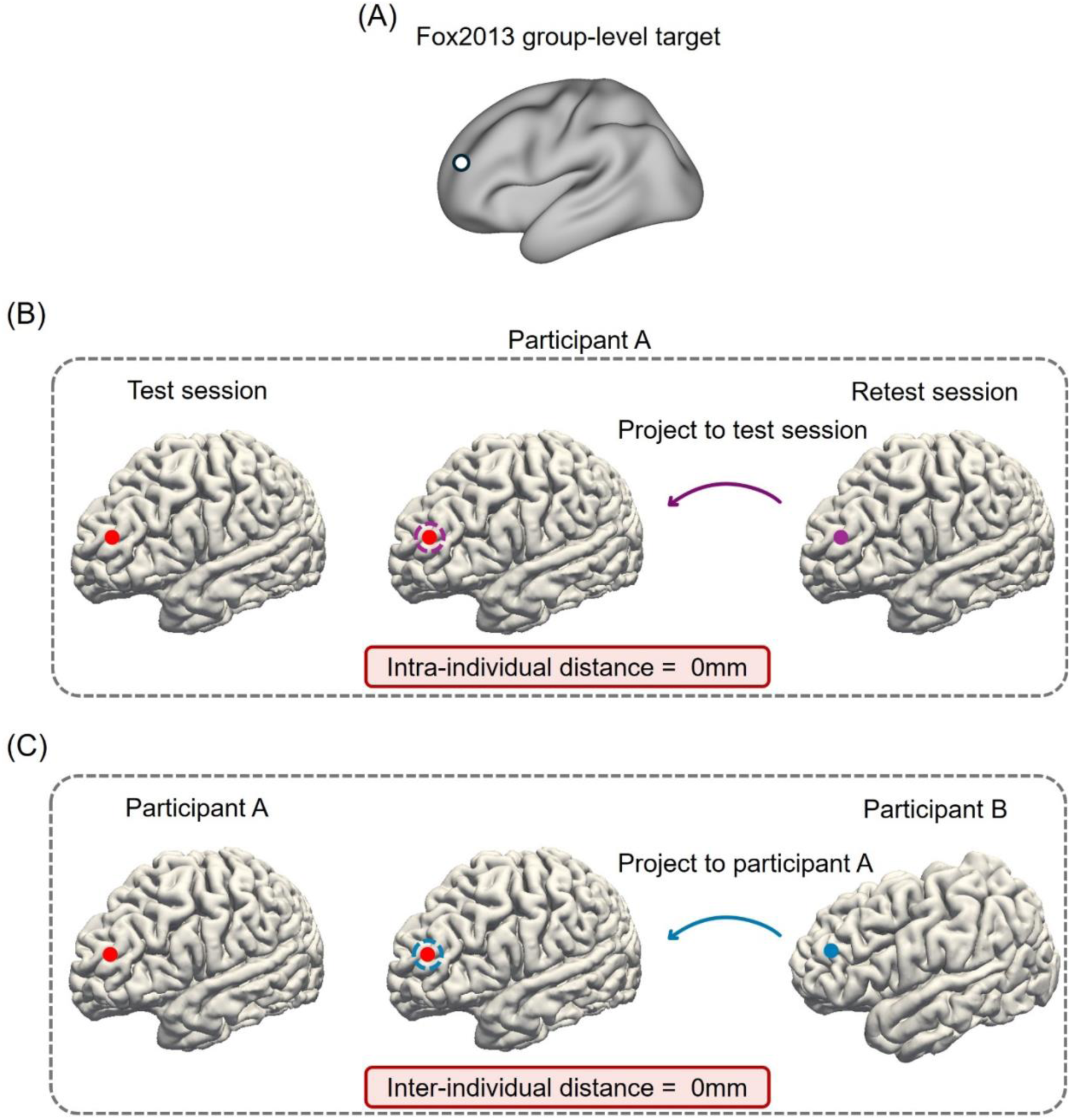
Illustration of a group-level target with perfect intra-individual reliability but no individual specificity. (A) A group-level target from MNI152 space (e.g., Fox et al., 2013). (B) The group-level target from MNI152 space (e.g., Fox et al., 2013) was anatomically projected to a participant’s T1 in the “test” session and the participant’s T1 in the “retest” session for neuromodulation. To compute test-retest reliability, the target in the “retest” session is projected to the “test” session (via MNI152 space) and compared with the target in the “test” session. In this scenario, the overlap between the transformed “retest” target and the “test” target will be perfect and the intra-individual distance will be zero. (C) On the other hand, suppose the group-level target from MNI152 space (e.g., Fox et al., 2013) was anatomically projected to participant 1’s T1 and ∂participant 2’s T1 for neuromodulation. To compute inter-individual variability, the target in participant 2’s T1 is projected to participant 1 T1 (via MNI152 space) and compared with participant 1’s target. In this scenario, the overlap between the transformed participant 2’s target and participant 1’s target will be perfect and the inter-individual distance will also be zero. Thus, there is perfect intra-individual agreement, but no individual specificity.

**Figure S2.**
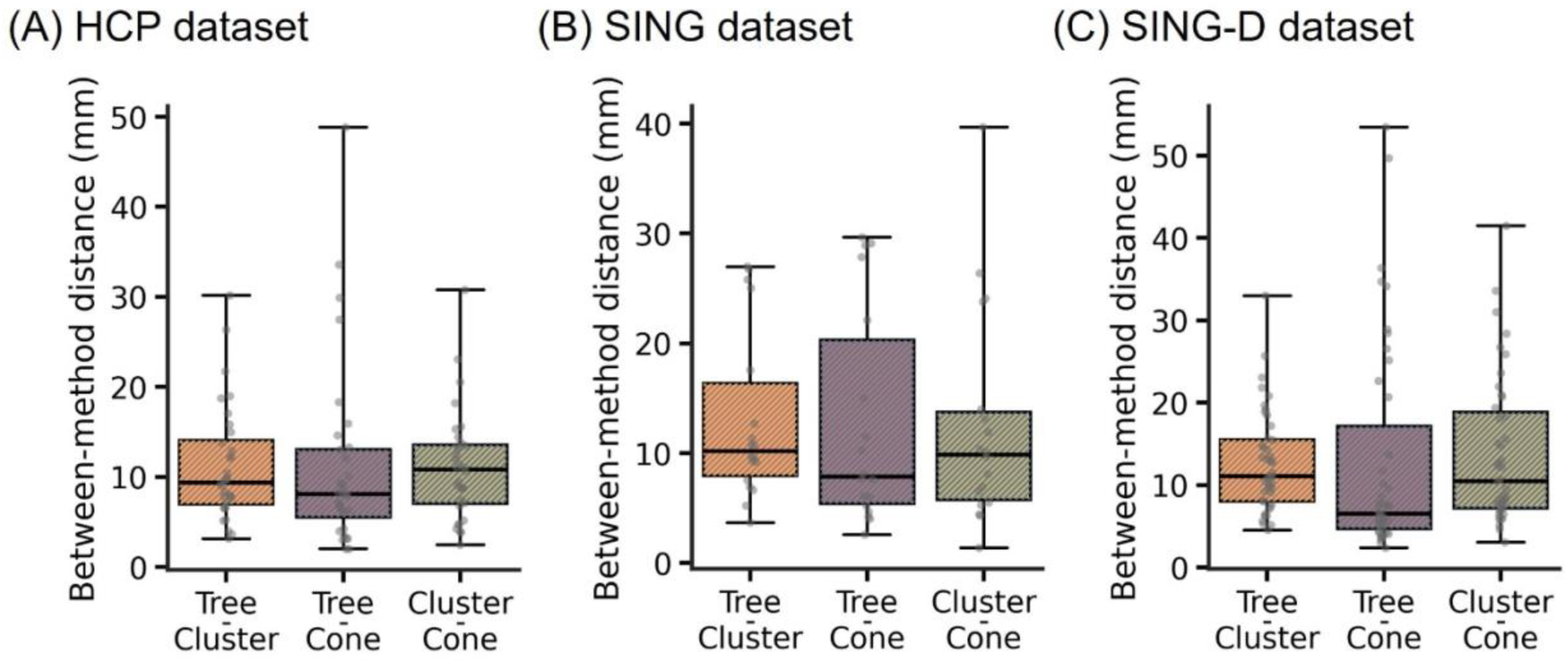
Pairwise Euclidean distances between depression targets derived from tree-based MS-HBM, cluster, and cone algorithms in (A) the HCP dataset, (B) the SING dataset, and (C) the SING-D dataset.

**Figure S3.**
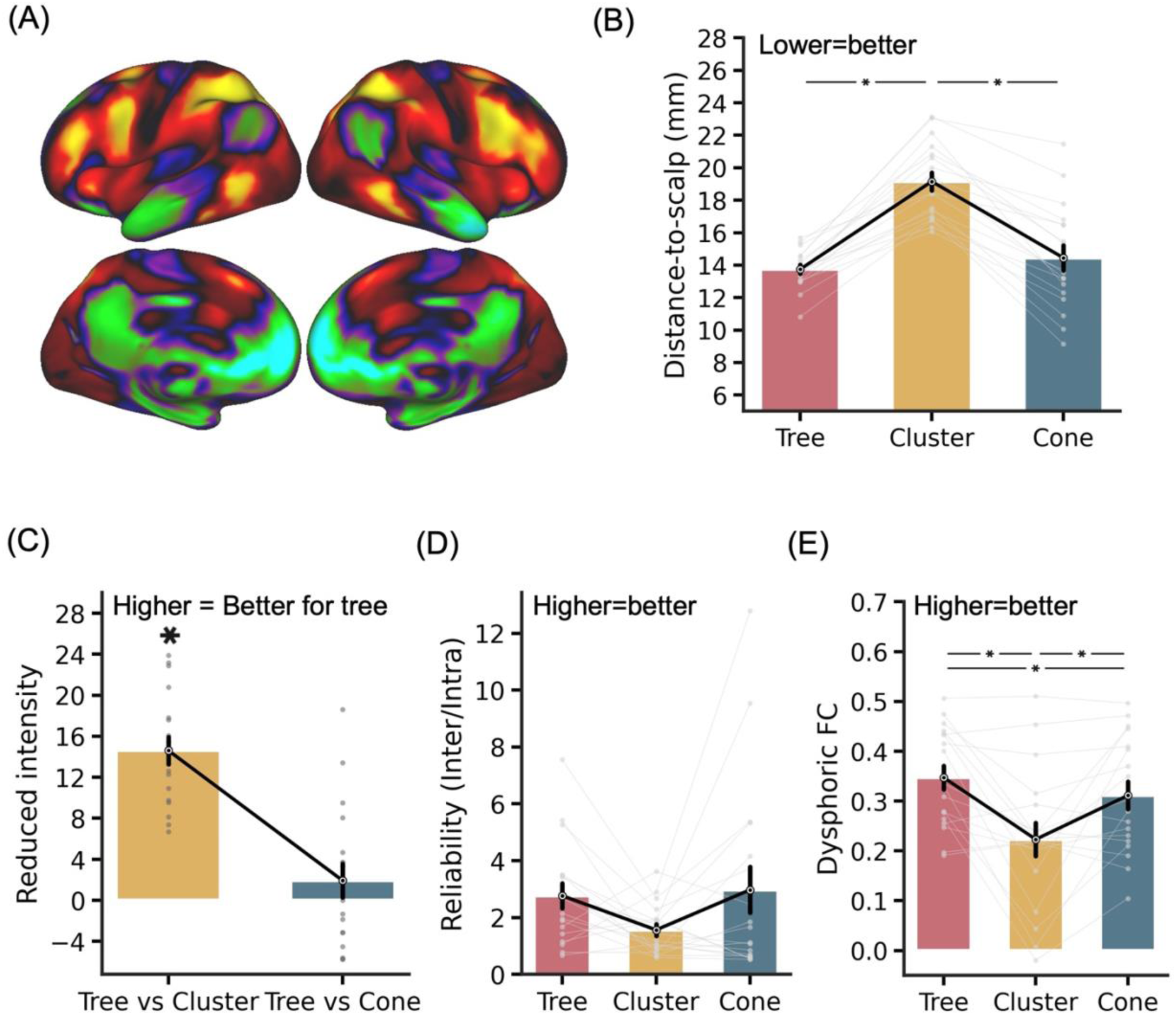
Personalized depression targets derived from tree-based MS-HBM using dysphoric circuit map. (A) Group-level dysphoric circuit map visualized on fsaverage6 surface. (B) Distance-to-scalp in the SING dataset. A smaller value indicates better performance. (C) Reduction in stimulation intensity in the SING dataset. Following the SAINT protocol (Cole et al., 2020), by using a linear adjustment in stimulation intensity based on distance-to-scalp and assuming 90% RMT dosage, a hypothetical reduction in stimulation intensity between tree-based MS-HBM and other approaches can be computed. (D) Reliability (ratio of inter-individual distance and intra-individual distance) in the multi-echo multi-band dataset. A higher value indicates better performance. (E) FC with the dysphoric circuit map in the SING dataset. A more positive value indicates better performance. * indicates statistical significance after multiple comparisons correction with FDR q < 0.05. We note that in all analyses, targets were derived in one session and then evaluated in another session.

## Footnotes

1 We note that commercial neuro-navigation softwares typically provide in-built proprietary registration algorithms that transfer targets from MNI152 space to individual native space for TMS. However, the neuro-navigation registration algorithms are typically different from those used in standard MRI preprocessing. Using one algorithm (e.g., ANTs) to register individual data to MNI152 space, computing the personalized target in MNI152 space and then using a different registration algorithm (from the neuro-navigation software) to project the target back to individual native space will introduce a potentially significant registration error.

2 The dosage formula is given by 0.9 **×** (RMT + 3 **×** (distance-to-scalp(target) - distance-to-scalp(M1))). Since the distance-to-scalp(M1) is the same across tree-based MS-HBM, cone and cluster algorithms, every mm decrease in distance-to-scalp(target) translates to 0.9 **×** 0.3 = 2.7% reduction.

